# Somatostatin interneurons in auditory cortex contribute to perceptual categorization

**DOI:** 10.1101/2022.07.06.498950

**Authors:** Ruiming Chai, Yuan Zhang, Yu Xin, Li Deng, Ning-long Xu

**Author notes:** Correspondence: Ning-long Xu.

## Abstract

In the mammalian neocortex, somatostatin (SOM)-expressing GABAergic interneurons serve as a critical circuit component, receiving diverse inputs and targeting various types of cortical neurons. However, their role in perception and cognition remains not well understood. In this study, we employed cell-type specific imaging and perturbation during an auditory perceptual decision task to investigate the contribution of SOM interneurons. Two-photon calcium imaging revealed that auditory cortex (ACx) SOM interneurons exhibit not only frequency-selective responses but also selectivity to categorical choices. Simultaneous optogenetic inactivation and two-photon imaging revealed that SOM interneurons refine the frequency discriminability of auditory cortical neurons and contribute to their selectivity to the categorical decision boundary. Moreover, bilateral chemogenetic inactivation of ACx SOM interneurons impaired behavior-level perceptual categorization. Our findings provide a mechanistic understanding on the functional significance of SOM interneurons in auditory cortical processing and perceptual categorization, linking cell-type specific cortical circuit computations with perceptual functions.

## Introduction

The mammalian neocortex contains diverse classes of inhibitory interneurons, which form distinct circuit motifs and exhibit various functional properties^1–4^. Three genetically distinct subtypes of GABAergic interneurons expressing parvalbumin (PV), vasoactive intestinal peptide (VIP) and somatostatin (SOM), respectively, constitute the majority of cortical inhibitory interneurons. Although the molecular, anatomical and physiological properties of these different subtypes of interneurons have been extensively characterized in the past decades, it was not until recently that the impact of specific subtypes of interneurons on cortical circuit processing and behavioral functions began to be unraveled^2^.

Significant progress has been made in understanding the circuitry and behavioral functions of PV interneurons^5–10^ and VIP interneurons^7,11–13^. The functions of SOM interneurons in behavior and cognition, however, remain less clear^14^. SOM interneurons possess unique functional properties, as they target the dendrites of pyramidal neurons^15–18^, and provide potent inhibition to other GABAergic interneurons and local excitatory neurons^19,20^. Additionally, SOM interneurons receive top-down feedback^21^ and neuromodulatory inputs^22^, and are modulated by sensory experience, behavioral state, and learning^23–26^. Similar to PV interneurons, SOM interneurons also contribute to cortical rhythmic activity^28,29^. Recent optogenetic manipulation studies show that SOM interneurons exert either a subtractive or divisive effect on sensory tuning properties of sensory cortical neurons^6,30–36^, and may participate in certain behavior-level functions^27,37^. Despite these functional implications, the precise role of SOM interneurons in regulating cortical circuit computations underlying specific perceptual and cognitive functions remains to be determined^2^. Our previous study showed that mice were able to perform auditory categorization on tone frequencies based on task rules^38,39^, during which auditory cortical neurons exhibited functional reorganization, implementing stimulus to category transformation^39^. We therefore asked whether SOM interneurons could contribute to the cortical computation for stimulus categorization.

Using cell-type specific in vivo two-photon imaging, we recorded SOM interneuron activity in the mouse auditory cortex during an auditory categorization task to assess task-relevant information coding in SOM interneurons. Furthermore, using an all-optical technique, we simultaneously imaged cortical population activity while optogenetically manipulating SOM interneurons in a trial-by-trial basis during task performance, which revealed the contribution of SOM interneurons to cortical representation for sensory stimuli and behavior-relevant categories. Finally, using chemogenetic inactivation, we investigated the causal contribution of SOM interneurons to categorical perceptual decision-making. Our results unravel crucial functions of SOM interneurons in cortical circuit computation for auditory perceptual categorization.

## Results

### Two-photon imaging of ACx SOM interneurons during auditory perceptual categorization

We trained head-fixed mice to perform an auditory-guided two-alternative-forced-choice (2AFC) task classifying various auditory tones into lower or higher frequency categories, and reporting their decisions by licking the left or right water spout^38,39^ (**Figure 1A**). Mice first learned to discriminate two pure tones with 2 octaves apart (8 and 32 kHz, 7 and 28 kHz or 5 and 20 kHz, stimulus duration, 300 ms). After reaching the criterion performance level (correct rate >85%), 4-6 additional tones at intermediate frequencies equally spaced in octaves between the two training tones were presented in randomly interleaved trials to test the categorization performance (**Figure 1A**; **STAR Methods**). To image SOM interneuron activity during task performance, we expressed the genetically encoded calcium sensor GCaMP6s^40^ in a Cre-dependent manner by injecting AAV-Syn-FLEX-GCaMP6s to unilateral auditory cortex of SOM-Cre mice, and implanted a chronic imaging window above the injection site. Ca^2+^ signals were recorded from 846 SOM interneurons in the layer 2/3 (L2/3) of auditory cortex using an *in vivo* two-photon microscope with a tiltable objective during the auditory decision-making task^39^ (**Figures 1B** and **1C**). Among all the imaged SOM interneurons, there are 574 neurons showing significant responsiveness during task performance (STAR Methods).

**Figure 1.**
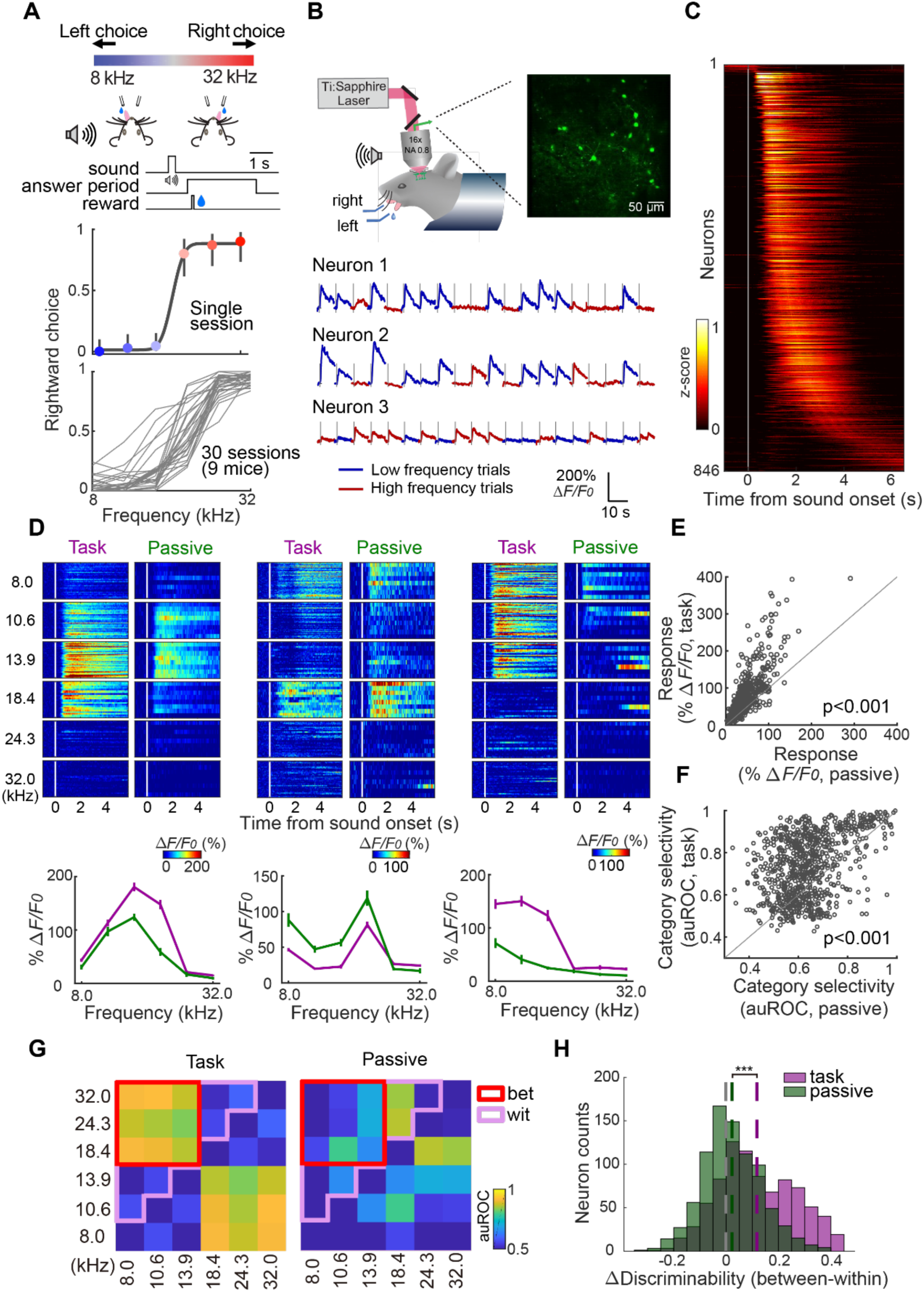
Two-photon calcium imaging of SOM interneurons in auditory cortex in behaving mice. (A) Auditory-guided two-alternative-forced-choice (2AFC) task. Top, schematic showing behavioral task configuration. Middle, psychometric curve of one example session. Error bar indicates 95% confidence interval. Bottom, task performances of all sessions. (B) Two-photon calcium imaging from auditory cortical SOM interneurons in behaving mice. Upper, experiment configuration with an example imaging field showing GCaMP expressing SOM interneurons in auditory cortex. Bottom, calcium traces from three example SOM interneurons recorded during task performance. The gray lines indicate sound onset time of each trial. (C) Calcium signals of all imaged SOM interneurons in task. Each row is Z-scored calcium signals averaged across trials from one neuron. The white line is sound onset time (n = 846 neurons, from 30 sessions, 9 mice). (D) Activity in task and passive state from three example SOM interneurons. Left, a neuron showing stronger responses in task than in passive state. Middle, a neuron showing weaker responses in task than in passive state. Right, a neuron showing higher category discriminability in task than in passive state. Upper, color raster plot of calcium signals in individual trials grouped by tone frequency. Lower, mean response amplitude as a function of frequencies. Purple, task; green, passive. Calcium signals are all aligned to sound onset time, indicated by the white vertical lines. Error bars indicate SEM. (E) Comparison of response amplitude in task and passive state for all imaged SOM interneurons. Each circle is the mean response amplitude from a single neuron (*P* = 3.31 × 10^−103^, Wilcoxon signed-rank test, n = 846 neurons, from 30 sessions, 9 mice). (F) Comparison of category discrimination (ROC analysis) in task and passive state. Each circle is a neuron (*P* = 4.68 × 10^−55^, Wilcoxon signed-rank test, n = 846 neurons, from 30 sessions, 9 mice). (G) Pairwise discriminability (auROC) of all frequencies from an example SOM interneuron, in task (left) and passive (right) state. Wit, within the same (low or high) frequency category; Bet, between low and high frequency categories. (H) Histograms showing distributions of the difference between ‘between-category’ and ‘within-category’ stimulus discriminability under task and passive conditions. The difference is significantly greater in task than that under passive condition (***, *P*=3.73 × 10^−53^, Wilcoxon signed-rank test, n = 846 neurons, from 30 sessions, 9 mice). Colored dash lines indicate the mean value of each distribution.

To assess the behavior-relevance of SOM interneuron activity, we recorded the Ca^2+^ signals during both task performance and passive listening while presenting the same set of tone stimuli (STAR Methods). We observed that ACx SOM interneurons exhibited clear sensory selectivity for different tone frequencies during both task performance and passive listening. While different neurons showed either enhanced or suppressed sound responses during task performance compared to passive listening (**Figures 1D**), the overall predominant effect was an enhancement of responses during task performance (**Figure 1E**; *P* =3.31 × 10^−103^, Wilcoxon signed-rank test). A subset of SOM interneurons showed selectivity to stimulus categories (corresponding to the two choices) rather than to single frequencies during task performance. Intriguingly, these neurons only show frequency selectivity but not category selectivity during passive listening (**Figure 1D, right**). Across the population, ACx SOM interneurons show significantly greater selectivity to category during task performance than during passive listening, as measured by receiver operating characteristic (ROC) analysis for each neuron (STAR Method; **Figure 1F**; *P* =4.68 × 10^−55^, Wilcoxon signed-rank test).

Our previous study showed that the category selectivity in ACx neurons manifested at the level of sensory coding during task performance. This was evidenced by the greater single-neuron discrimination for pairs of stimuli from different categories compared to stimuli from the same category^39^. Here, we examined whether the frequency discrimination by SOM interneurons also exhibits category related modulation. Single neuron discrimination between individual stimuli based on ROC analysis showed that under the task condition, discrimination was significantly greater between stimuli from different categories than those from the same category (**Figures 1G and 1H**, *P* = 2.29×10^−76^, Wilcoxon signed-rank test, n = 846 neurons,). Such difference between within-and between-category stimulus discriminability was significantly greater under the task condition than that under the passive listening condition (**Figures 1G and 1H**, task auROC difference = 0.117±0.005; passive auROC difference = 0.023±0.004; *P* = 3.73×10^−53^ between task and passive conditions, Wilcoxon signed-rank test, n = 846 neurons).

These results indicate that ACx SOM interneurons are not only selective to sensory stimuli, but also show task-dependent category selectivity, both markedly enhanced by the engagement of behavioral task.

### Task modulation of SOM activity emerges after learning

The marked difference in SOM interneuron responses between task performance and passive listening states appears to reflect task-specific demands for perceptual categorization of auditory stimuli, consistent with our previous study^39^. However, there could be other general state differences between task performance and passive sensory stimulation conditions that were not directly pertinent to the sound categorization performance, for instance, the thirsty state, licking-related movements and reward-related processes. These general differences should be present when the animal has learned the task structure of the 2AFC task but is not yet proficient in the stimulus categorization. We thus compared the task-dependent modulation of SOM activity between early learning (after having learned the basic structure of the 2-choice task, but with categorization performance <60% correct rate) and well-trained stages (categorization performance >80% correct rate) by performing two-photon imaging from the same population of SOM interneurons across these learning stages. We tracked the activity of 88 ACx SOM interneurons across task learning (**Figures 2A** and **2B**), and examined the responses during task performance and passive listening both in early learning stage (correct rate < 60%) and in expert stage (correct rate > 80%). As shown in **Figures 2C** and **2D**, during the early task training stage, a SOM interneuron exhibited comparable sensory tuned responses in both task and passive conditions (**Figure 2C**). In the expert stage, this same interneuron displayed notable category selectivity during task performance, favoring the low-frequency tone category. However, in the passive state of the expert stage, the neuron still exhibited sensory-tuned responses akin to those observed during the early learning stage (**Figure 2D**). Across the population, SOM interneuron responses showed no significant differences between task and passive conditions in the early learning stage (**Figure 2E**). In contrast, the responses of the same group of neurons were significantly enhanced by task performance in the expert stage (**Figure 2F**). Likewise, the same population of neurons only show significant task enhancement of coding for stimulus categories in the expert stage (**Figure 2H**), but not in the early learning stage (**Figure 2G**). These results suggest that the task-related enhancement of responses and the emergence of category coding in SOM interneurons were due to task-specific modulation after mice learned the task.

**Figure 2.**
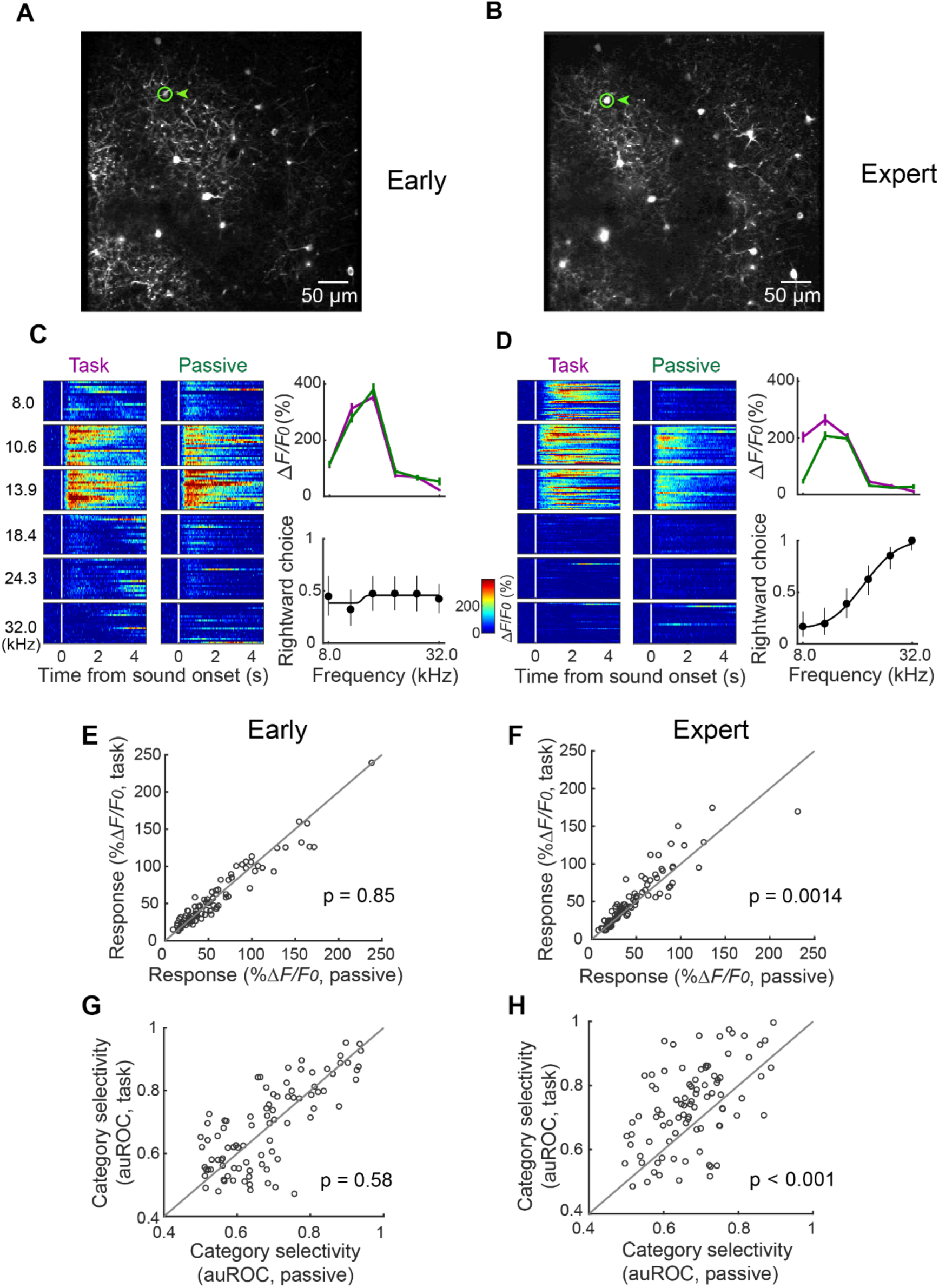
Task modulation of SOM interneurons emerged after task learning. (A) and (B) Two-photon images of SOM interneurons from one example field in early and expert stage respectively. Green circle and triangle indicate the example neuron. (C) Activity of the example neuron indicated in (A) in task and passive states in early learning stage. Behavioral performance is shown as the psychometric curve on the right. (D) As in (C) showing the activity from the same neuron and the behavioral performance in expert stage. (E) Comparison of response amplitude of individual neurons under task and passive conditions in early learning stage (*P* = 0.85, Wilcoxon signed-rank test; n = 88 neurons from 5 imaging fields, 5 mice). (F) Same as (E), but in expert stage (*P* = 0.0014, Wilcoxon signed-rank test, n = 88 neurons from 5 imaging fields, 5 mice). (G) Comparison of category selectivity of individual neurons under task and passive conditions in early learning stage (*P* = 0.58, Wilcoxon signed-rank test; n = 88 neurons from 5 imaging fields, 5 mice). (H) Same as (G), but in expert stage (*P* = 1.01 × 10^−5^, Wilcoxon signed-rank test, n = 88 neurons from 5 imaging fields, 5 mice).

### Encoding of different task variables by individual SOM interneurons

We next ask how the activity of SOM interneurons is modulated by different task variables, including sensory, choice, reward and licking movement. Our 2AFC task with multiple tone frequencies provides a variety of trial types in which different task variables could be decorrelated. We thus used a generalized linear model (GLM) to quantify the simultaneous contribution of these task variables to the activity of individual SOM interneurons (**Figure 3A, STAR Methods**)^41^. We found that for most SOM interneurons, tone frequency and category choice provided the primary contributions, with little contributions from reward and lick number (**Figure 3B**). We next sorted the neurons based on the dominant contributing variable of each neuron (**Figure 3C**). Frequency dominant and choice dominant neurons constituted two primary and distinct clusters, suggesting that sensory and choice information are represented by different SOM interneuron populations. We further separately examined the relative contributions of different variables to frequency coding and choice coding neurons. For SOM interneurons showing predominant sensory coding (frequency contribution > 0.3), choice contributions were significantly weaker (**Figure 3D**). Similarly, tone frequency provided significantly weaker contributions for neurons with prominent choice coding (choice contribution > 0.3, **Figure 3E**). In both choice and sensory coding SOM interneurons, the contributions from reward and lick number were minimal (**Figure 3D and 3E**).

**Figure 3.**
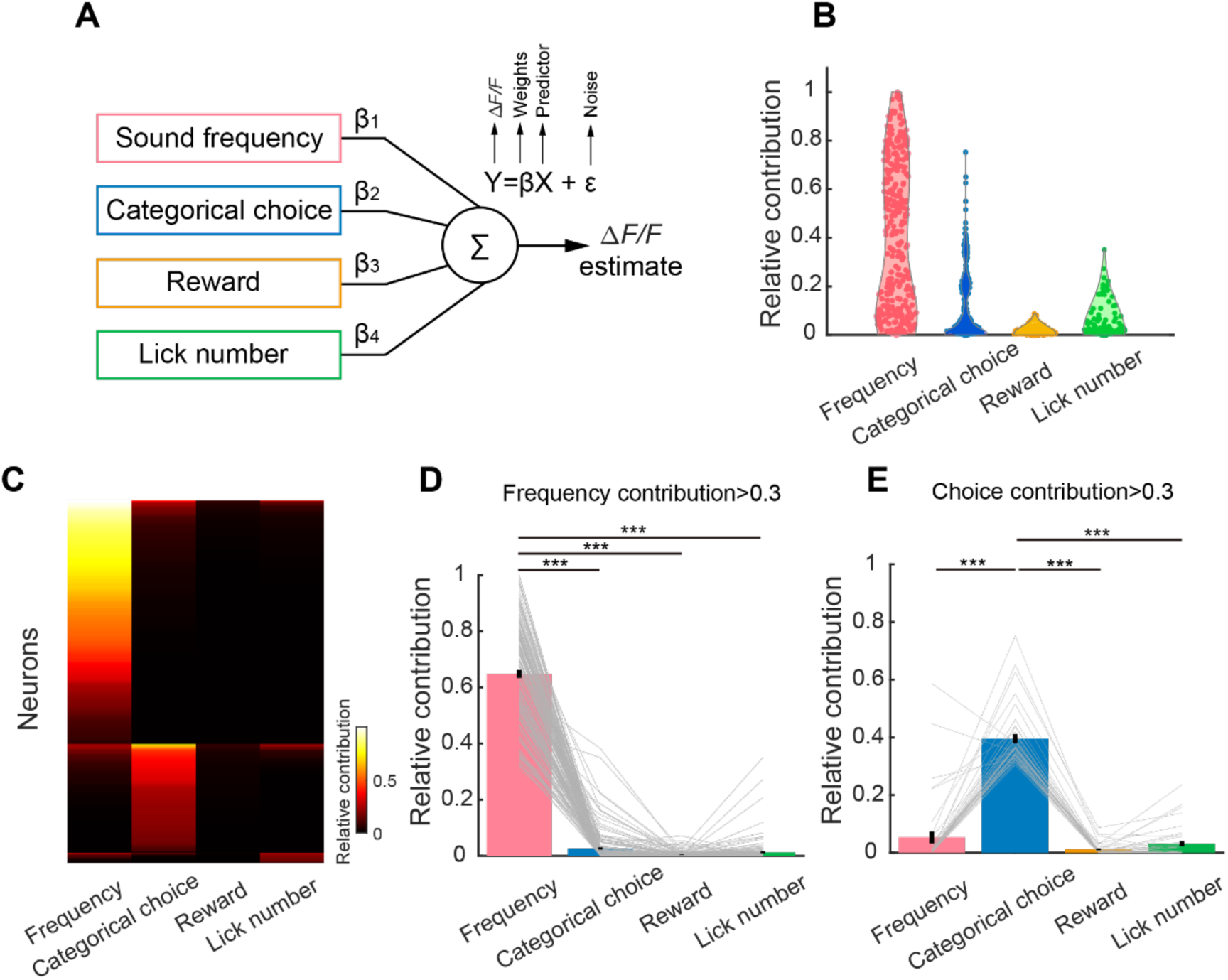
Modeling the contributions of behavioral variables to SOM interneuron activity using a generalized linear model (GLM). (A) Schematic of the GLM model using behavioral variables to predict SOM neuron activity. (B) Relative contributions to explained variance of neuronal activity by different behavioral variables, for SOM interneurons with significant amount of activity variance explained by the model (see STAR Methods, n = 350 neurons). (C) Heatmap representing relative contributions by different behavioral variables to each SOM interneuron activity. Neurons were first grouped by the variable that shows the maximal contribution, and then sorted by the contribution value. (D) Comparison of relative contributions from different variables for SOM neurons with prominent sensory coding (frequency contribution > 0.3, n = 175; ***, *P* < 0.001, Wilcoxon signed-rank test). (E) Comparison of relative contributions from different variables for SOM neurons with prominent choice coding (choice contribution > 0.3, n = 40; ***, *P* < 0.001, Wilcoxon signed-rank test).

### Simultaneous optogenetic inhibition of SOM interneurons and two-photon population imaging

To investigate the role of SOM interneurons in regulating cortical circuit computations during auditory perceptual decision-making, we employed simultaneous *in vivo* two-photon imaging and optogenetics to record local population activity while inactivating SOM interneurons during task performance (**Figure 4A**). We injected a mixture of AAV-hSyn-FLEX-Jaws-tdTomato and AAV-hSyn-GCaMP6s in L2/3 of the auditory cortex of SOM-Cre mice. This allowed Jaws^42^ (a red light activated activity suppressor) to be expressed only in SOM interneurons in a Cre-dependent manner, and GCaMP6s was expressed ubiquitously in the L2/3 neurons (**Figure 4A**). Red light (635 nm) was delivered to the imaging field through the objective of the two-photon microscope equipped with a separate stimulating light path via a multi-dichroic filter to inhibit SOM interneurons expressing Jaws^43^ (**Figure 4A**, see **STAR Methods**). Calcium signals from neurons in the same region, indicated by GCaMP6s, were acquired simultaneously using two-photon microscopy. Photostimulation of Jaws using red light was presented in about half of the trials, which were randomly interleaved with control trials (no red light stimulation). Photosimulation strongly inhibited the responses of SOM interneurons expressing Jaws (**Figure 4B**-**4E**). But for other cortical neurons, which were putatively non-SOM neurons, red light stimulation led to an increase in the response amplitude in majority of these neurons (**Figures 4F-4H**), indicating a general inhibitory effect of SOM interneurons on the local cortical population.

**Figure 4.**
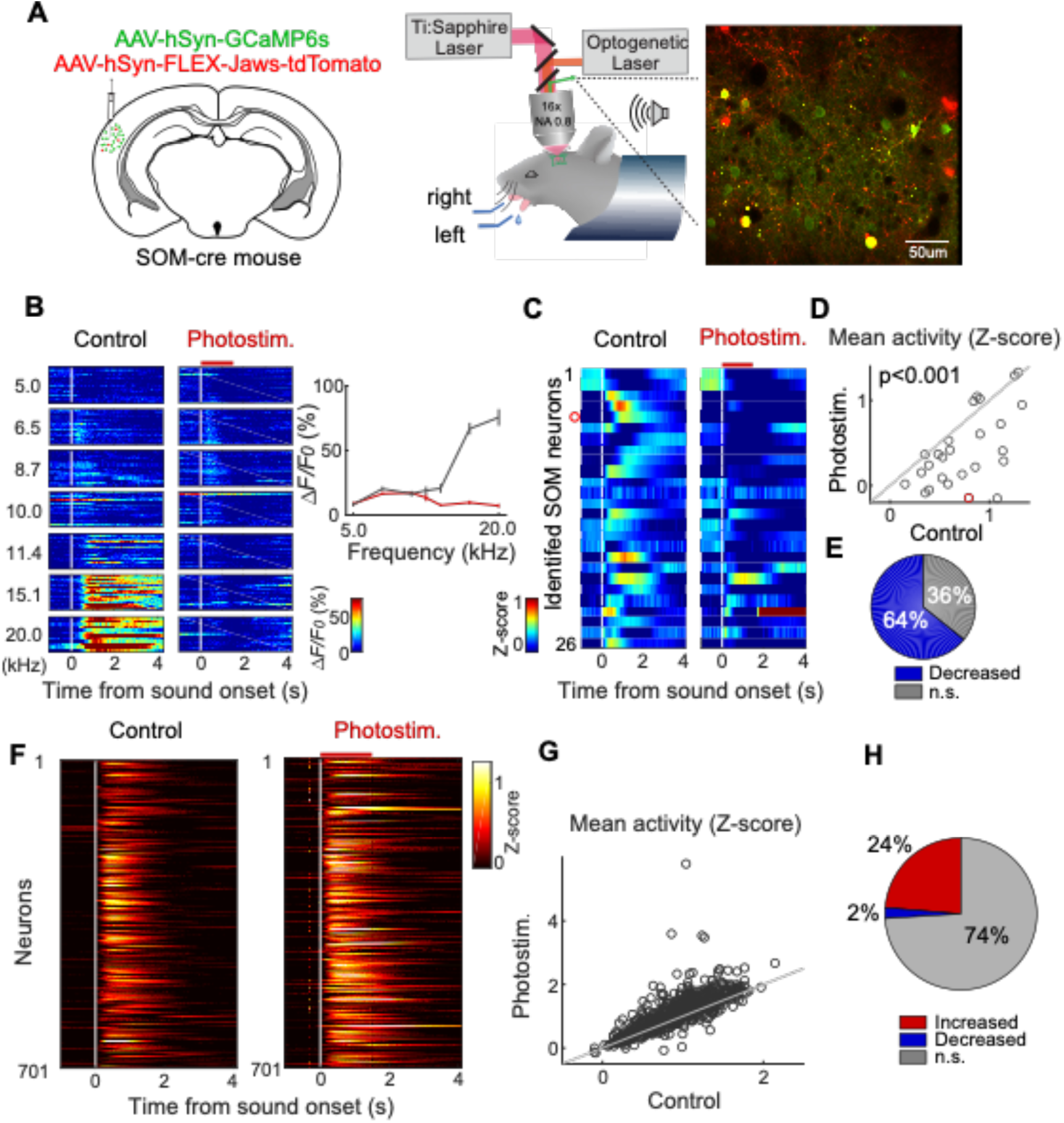
Simultaneous optogenetic inhibition of SOM interneurons and two-photon imaging. (A) Schematic showing the virus injection (left) and simultaneous two-photon calcium imaging of auditory cortical neurons and optogenetic manipulation of SOM interneurons (right). Inset: one example imaging field. Green, neurons expressing GCaMP6s; red, SOM interneurons expressing Jaws; yellow, SOM interneurons expressing both GCaMP6s and Jaws. (B) Activity of one example identified SOM interneuron co-expressing Jaws and GCaMP6s, under control and photostimulation conditions. The red horizontal line indicates photostimulation time. (C) Z-scored activity of all identified SOM interneurons under control and photostimulation conditions. Each row is the Z-scored activity averaged across all trials from one neuron (n = 26 neurons, from 7 sessions, 3 mice). The red circle indicates the example neuron in (B). (D) Comparison of response amplitude under control and photostimulation conditions for all identified SOM interneurons. Each circle is the Z-scored peak response amplitude in 1 s time window after sound, averaged across all trials from one neuron (*P* = 1.62 × 10^−4^, Wilcoxon signed-rank test, n = 26 neurons, from 7 sessions, 3 mice). The red circle indicates the example neuron in (B). (E) Quantification of activity change induced by photostimulation manipulation of all identified SOM interneurons using ANOVA (n = 26 neurons, from 7 sessions, 3 mice). (F) Calcium signals of all imaged putative non-SOM neurons in task with (right) or without (left) photostimulation. Each row is Z-scored calcium signals averaged across trials from one neuron. The white vertical line is sound onset time. The red horizontal line indicates photostimulation time (n = 701 neurons from 7 sessions, 3 mice). (G) Response amplitudes in photostimulation trials plotted against those in control trials for all imaged putative non-SOM neurons. Each circle is the Z-scored response amplitude from a single neuron (n = 701 neurons from 7 sessions, 3 mice). (H) Quantification of activity change induced by photostimulation manipulation of all putatively non-SOM neurons using ANOVA (n = 701 neurons, from 7 sessions, 3 mice).

### SOM interneurons contribute to stimulus discrimination and categorization in ACx local circuits

In addition to the general inhibitory effect, SOM interneurons may have more precise influences on local circuit processing. To investigate this, we analyzed the information coding of individual ACx neurons during task performance with or without photoinhibition of SOM interneurons. To understand the contributions of SOM interneurons to sensory discrimination, we first examined auditory selective but not choice selective neurons (**STAR Methods**). Closer inspection revealed structured changes in the response properties of local cortical neurons following SOM interneuron photoinhibition. In one subset of neurons (type I), SOM interneuron silencing resulted in a more pronounced enhancement of responses to non-preferred stimuli than to preferred stimuli, yielding a broadened tuning width (**Figures 5A** and **5C**). This implies that under normal conditions, SOM interneurons exert selective suppression to activity other than the preferred frequency evoked responses, and consequently sharpen the frequency tuning of these neurons. In another subset of neurons (type II), photoinhibition of SOM interneurons preferentially amplified responses to the preferred frequency, indicating that SOM interneurons normally exert a suppressive scaling effect on the stimulus tuning of these neurons (**Figures 5B**). While in the example type I neuron, the sharpening effect appears to be asymmetrical for the frequencies flanking the preferred frequency (**Figure 5A**), summarized tuning curves of type I neurons exhibits a symmetrical broadening effect following photoinhibition of SOM interneurons (**Figure 5C**), suggestion that SOM interneurons normally sharpen the sensory tuning of the type I population. In contrast, for type II neurons, the population tuning curve was not changed after photoinhibition of SOM interneurons (**Figure 5F**). Instead, when summarizing the responses of type II neurons as a function of stimulus ranked by response amplitude, the scaling effect is manifested as the stronger inhibition by SOM interneurons for stimuli closer to the best frequency (**Figure 5G**), a trend not present in the type I neurons (**Figure 5D**).

**Figure 5.**
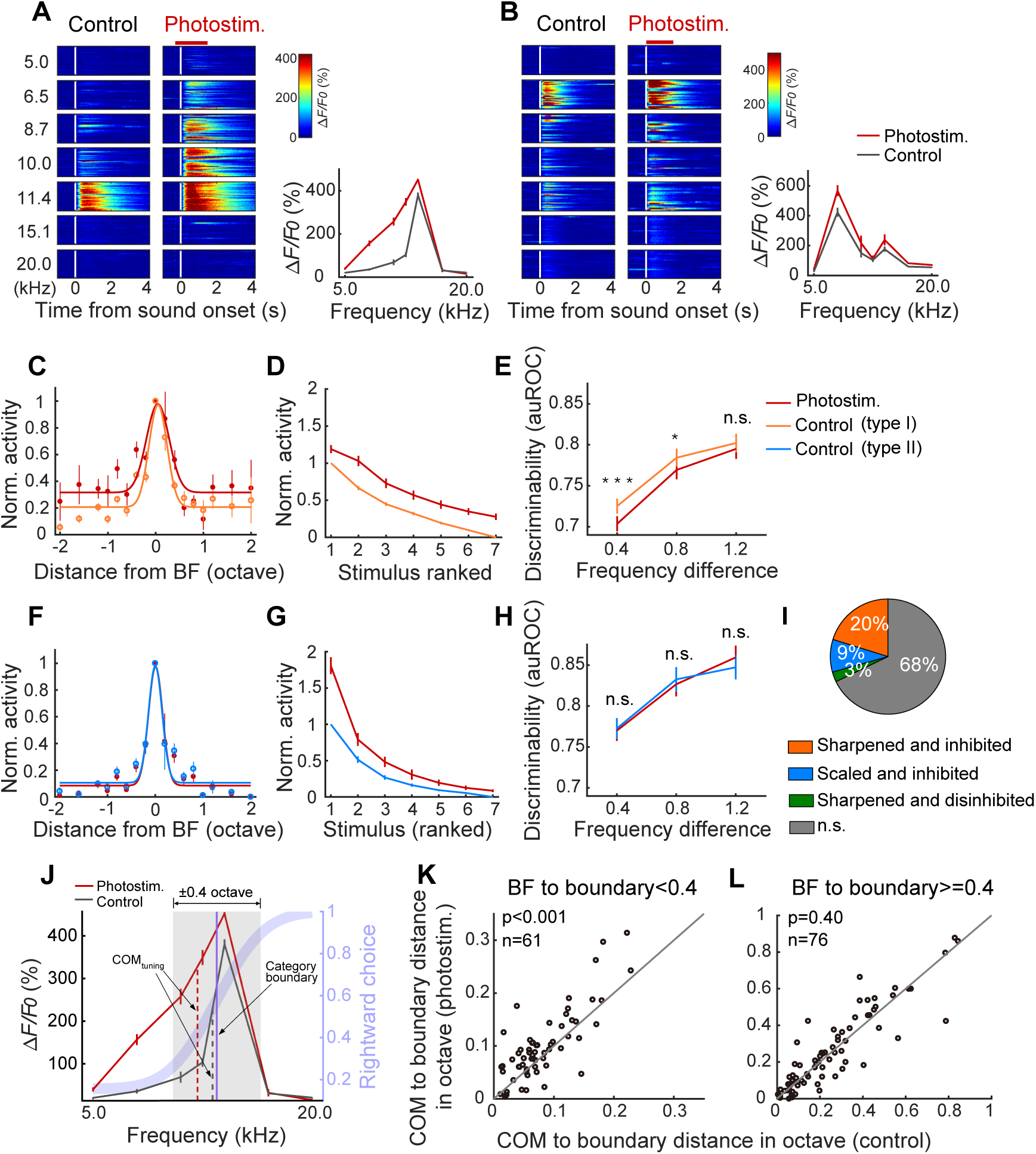
Contribution of SOM interneurons to sensory representations in ACx local circuits. (A) Activity of an example neuron sharpened by SOM interneurons. Left, color raster plot of calcium signals in individual trials grouped by tone frequency, with or without photoinhibition of SOM interneurons. The red horizontal line indicates photostimulation time. Right, tuning curves under control and photostimulation conditions. Black, control condition; red, photostimulation condition. Error bars indicate SEM. (B) Same as (A), for an example neuron exhibiting scaling effect by SOM interneurons. (C) Normalized responses as a function of tone frequency offset to BF for all type I neurons (sharpened by SOM interneurons). The responses were normalized to the range between 0 and 1, by firstly subtracting the minimal response and then dividing them by the difference between BF response and the minimal response of the corresponding condition. Orange, control condition; red, photostimulation condition. Error bars indicate SEM (n = 98 neurons, from 7 sessions, 3 mice). (D) Normalized responses ranked by frequency preference and then averaged across neurons, of all type I neurons. Responses under both conditions were both normalized to control conditions, by firstly subtracting the minimal response (control condition) and then dividing them by the difference between BF response (control condition) and the minimal response (control condition). Orange, control condition; red, photostimulation condition (n = 98 neurons, from 7 sessions, 3 mice). (E) Discriminability of tone stimulus pairs with various frequency differences, for each single type I neuron under control condition (orange) and photostimulation condition (red). Discriminability (auROC) is grouped and sorted by octave distance of each tone pair, averaged across tone pairs within the same group, and then averaged across neurons. Error bar indicates SEM (n = 98 neurons, from 7 sessions, 3 mice. n.s., no significance; *, *P* < 0.05; ***, *P* < 0.001, Wilcoxon signed-rank test). (F)-(H) Same as (C)-(E), for type II neurons (scaled by SOM interneurons). Blue, control condition; red, photostimulation condition (n = 39 neurons, from 7 sessions, 3 mice). (I) Fraction of neurons of showing different types of modulation follwing SOM interneurons, among all sound responsive neurons (n = 427 neurons from 7 sessions, 3 mice). (J) Tuning curves of the example neuron in (A), under control (black) and photostimulation (red) conditions. Vertical dashed lines indicate center of mass of tuning curves (COM_tuning_) under the corresponding condition. Purple curve: the behavioral psychometric function. Purple vertical line: category boundary estimated from the psychometric function. Gray shadow indicates ±0.4 octave range centered around the behavioral category boundary. Error bars indicate SEM. (K) Distance from COM_tuning_ to the behavioral category boundary, compared between control and photostimulation conditions for neurons with BF near the category boundary (distance < 0.4 octave). (L) Same as (K), but for neurons with BF far from the category boundary (distance >= 0.4 octave).

The sharpening of frequency tuning (type I) is likely to increase neuronal stimulus discriminability, while the scaling of the tuning curve (type II) is likely to influence stimulus detection. Since the behavioral performance in our task primarily depends on stimulus discrimination rather than detection^39^ (**Methods**), the type I modulation may play a more important role. We thus asked how SOM interneurons contribute to stimulus discrimination by individual neurons as measured using ROC analysis. Indeed, for type I neurons, silencing SOM interneurons significantly reduced neuronal discrimination for stimulus pairs with smaller frequency differences (**Figure 5E**), suggesting that SOM interneurons contribute to the fine discrimination of stimulus by these neurons. For type II neurons, however, silencing SOM interneurons did not significantly change stimulus discriminability (**Figure 5H**). Notably, there are greater proportion of type I neurons (23%) than type II neurons (9%) that were recorded during task performance (**Figure 5I**), suggesting that SOM interneurons more prominently contribute to accurate discrimination of stimulus by local population, paralleling with the task demand.

The current behavioral task requires not only stimulus discrimination but also stimulus categorization. We wondered whether SOM interneurons may contribute to the cortical processing for stimulus categorization. Stimulus categorization comprise two processing stages: 1) the computation of stimulus to category transformation and 2) category to motor choice transformation. We first examined whether SOM interneuron could contribute to categorical choices (**Figure S5**). We identified choice selective L2/3 ACx neurons (**Figure S5A-S5F**) and examined their activity changes following SOM inactivation. We found that while SOM interneurons primarily exert a general inhibitory effect on choice selective neurons (**Figure S5G**), the selectivity for category choices remains unchanged following SOM inactivation (**Figure S5H**), suggesting that SOM interneurons do not directly contribute to the choice representation.

We next examine whether SOM interneurons may contribute to the stimulus to category transformation. Our previous study showed that the auditory cortical neurons contribute to stimulus to category transformation by enhancing the representations for stimuli that are close to the category boundary^39^. We therefore asked how photoinhibition of SOM interneurons may influence the representations for near-boundary stimuli. To address this, we separately examined the effect of SOM photoinhibition on neurons with higher and lower degrees of representation for the category boundary. To estimate the degree of category boundary representation, we examined the relationship between frequency tuning by measuring the distance from the center of mass of tuning curve (COM_tuning_) to the category boundary. We then asked how this distance may change following SOM inactivation (**Figure 5J**). For neurons showing higher degrees of representation for category boundary, with BF close to the category boundary (< 0.4 octave), we found that the COM_tuning_ to boundary distance significantly increased after SOM inactivation (**Figures 5J and 5K**). However, for neurons with lower degree of boundary representation (with BF >0.4 octave from the category boundary), the COM_tuning_ to boundary distance was not significantly changed after SOM inactivation (**Figure 5L**). These results indicate that SOM interneurons may contribute to the representations for near-boundary stimuli, and therefore participate in the computation of stimulus to category transformation.

### Causal contribution of SOM interneurons to perceptual discrimination

The above assessment at the neuronal level indicates that SOM interneurons contribute to the discrimination of more difficult stimulus pairs (difference <1 octave) by individual cortical neurons. We asked whether this effect would manifest at the behavioral level during decision-making task. To address this, we reversibly silenced SOM interneurons of the auditory cortex in both hemispheres during task performance using chemogenetics^44,45^. We expressed hM4Di in SOM interneurons in auditory cortex by bilateral injection of AAV-hSyn-FLEX-hM4Di-mCherry in SOM-Cre mice (**Figure 6A**). Clozapine-N-oxide (CNO) or saline (as control) was administered in the same animals via intraperitoneal (IP) injection prior to each behavioral session on alternating days. We found that compared to saline injection, inactivation of SOM interneurons in auditory cortex following injecting CNO significantly impaired behavioral performance of auditory discrimination as indicated by reduced slope of psychometric functions (**Figures 6B** and **6C**), suggesting an impaired level of stimulus categorization^38^. Consistent with this notion, when we separately examine the animals’ task performance in difficult trials (with frequencies closer to the discrimination boundary) and easy trials (with frequencies on the two ends of tested frequency range), we found that inactivation of SOM interneurons significantly reduced task performance only in difficult trials (**Figure 6D**), but not in easy trials (**Figure 6E**). The effects are consistent across animals (**Figure 6F-H**). These results suggest that SOM interneurons causally contribute to behavioral level stimulus categorization by enhancing discrimination of stimuli closer to the category boundary. This is consistent with the neuronal level causal contribution of SOM interneurons to local cortical stimulus representations (**Figure 5**). To control for any non-specific effect of CNO injection, we expressed tdTomato in SOM interneurons of auditory cortex by bilateral injection of AAV-hSyn-FLEX-tdTomato in SOM-Cre mice (**Figure 6I**), and found no significant changes in psychometric slopes or in the performance on difficult or easy trials following CNO injection (**Figures 6J** to **6L**). Together, these results indicate that SOM interneurons play a causal role in auditory perceptual decision-making by contributing to finer-level discrimination.

**Figure 6.**
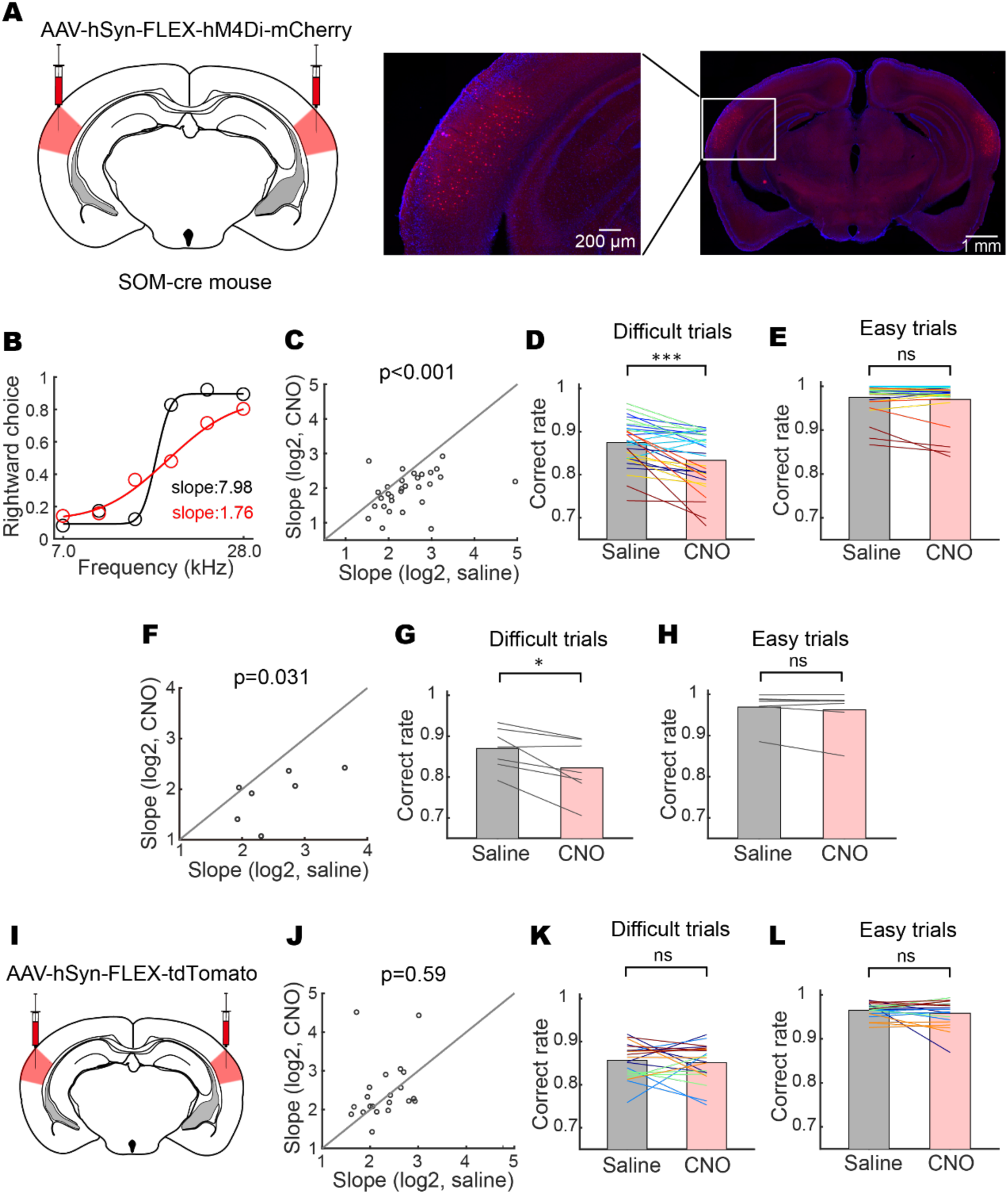
Inhibition of SOM interneurons impairs behavioral performance on perceptual discrimination. (A) Left, schematic showing the virus injection site expressing hM4Di in SOM interneurons. Right, histology imaging showing viral expression in auditory cortex. (B) Psychometric functions showing behavior performance from a CNO injection session (red) and a saline injection session (control, black). (C) Statistical comparison of psychometric function slope in saline and CNO sessions (*P* = 6.89×10^−5^, Wilcoxon signed-rank test, n = 30 session pairs from 7 mice, 3-5 session pairs from each mouse). (D) Statistical comparison of correct rate of difficult trials (intermediate frequencies between 7 and 28 kHz) in saline and CNO sessions. Lines are data from alternating sessions with saline or CNO. Different colors represent session pairs from different animals (*P* = 8.19×10^−5^, Wilcoxon signed-rank test, n = 30 session pairs from 7 mice). (E) Same as (D), but for easy trials with end frequencies, 7 and 28 kHz (*P* = 0.103, Wilcoxon signed-rank test, n = 30 session pairs from 7 mice). (F) – (H) Statistical comparison across animals for psychometric function slope (F), correct rate of difficult trials (G), correct rate of easy trials (H). Each data point is the data averaged across sessions of an individual animal. Wilcoxon signed-rank test, *P* = 0.031 (F), *P* = 0.031 (G), *P* = 0.47 (H), n = 7 mice. (I)-(L) Control experiments with tdTomato expressed in SOM interneurons in auditory cortex. Data are presented as in (C)-(E). Wilcoxon signed-rank test, *P* = 0.59 (J), *P* = 0.72 (K), *P* = 0.77 (L), n = 21 session pairs from 5 mice, 4 or 5 session pairs from each mouse.

## Discussions

To understand how behaviorally relevant computations are implemented in cortical circuits requires an appreciation of how distinct interneuron subtypes affect local networks during well-defined behavioral tasks. Earlier studies using *ex vivo* preparations revealed rich mechanisms for how different subtypes of interneurons affect local circuit properties^3,17,20,46–48^. In recent years, with the availability of genetic targeting tools^49,50^ and cell-type specific recording and manipulation methods^51,52^, the functions of specific subtypes of inhibitory interneurons have been extensively investigated in intact mouse brain under anesthetized or quiet awake condition^6,8,29,35,36,53–55^. But to understand the impact of specific subtypes of inhibitory interneurons on cortical circuit computations, it is necessary to address this problem under well-defined behavioral tasks that make use of such computations and assess its contribution to behavioral functions^2^. Here we directly examined the functional role of SOM interneurons at both local circuit level and at behavioral level during a perceptual discrimination task. We found that ACx SOM interneurons encode both auditory information and task-related categorical choice information (**Figure 1**), the latter of which emerged after the animals became proficient in the specific task (**Figure 2**). Simultaneous optogenetic manipulation and two-photon imaging revealed that ACx SOM interneurons exert either sharpening or scaling effect on the auditory representations of different groups of local L2/3 neurons during task performance. Interestingly, a greater proportion of the modulated neurons showed sharpened tuning by SOM interneurons (**Figure 5I**), a property more relevant to the current perceptual categorization task. Such effect on finer stimulus discrimination in local cortical neurons likely further contributes to the representation of category-boundary (**Figures 5J-5L**), which may facilitate the computation for stimulus categorization^39^. Consistent with this effect on local circuit processing, chemogenetic silencing of ACx SOM interneurons during task revealed that SOM interneurons specifically contribute to perceptual discrimination of more similar auditory stimuli, resulting in a steeper psychometric slope reflecting increased perceptual sensitivity (**Figure 6**). Together, these results unraveled a task-specific role of ACx SOM neurons in both cortical sensory processing and perceptual discrimination, linking cortical circuit computations to a well-defined behavioral function.

In accordance with prior studies on sensory evoked responses in SOM interneurons^18,54–57^, we observed that SOM interneurons in the auditory cortex exhibit reliable responses to sound stimuli during both passive stimulation and task performance (**Figure 1**), displaying a seemingly longer response onset compared to other types of cortical neurons (**Figures 1** and **4**). This observation aligns with the notion that SOM interneurons in the sensory cortex primarily receive inputs from cortical feedback connections rather than directly from ascending thalamic projections^56,57^. Intriguingly, we also observed a task-dependent enhancement of SOM responses (**Figure 1**), which may indicate top-down modulation from higher-order areas^21^ or from neuromodulatory inputs^22^.

When targeting excitatory neurons, SOM interneurons primarily innervate the apical dendrites of cortical pyramidal neurons, exerting more specific control over synaptic integration—a process that can be distinct from the direct inhibition of spiking output by PV interneurons targeting perisomatic domains. Furthermore, SOM interneurons also target other subtypes of interneurons, including both PV and VIP interneurons. As a result, instead of mediating conventional feedforward^54,56^ or feedback inhibition^17^, SOM interneurons may serve as a versatile regulator in local cortical circuits, regulating the integration of top-down behavioral state information and bottom-up sensory input during active perceptual behavior. The distinct nature of SOM interneurons necessitates a thorough investigation of their contributions to circuit and behavioral functions.

Employing simultaneous two-photon population imaging and cell-type specific optogenetic inactivation, we evaluated the contribution of SOM interneurons to cortical circuit computations during a behavioral task that specifically requires such computations. Our findings indicate that SOM interneurons either sharpen auditory frequency tuning or scale down the tuning peak of various groups of local cortical neurons (**Figure 5**). A previous study in the auditory cortex of mice also reported similar dual effects on sensory tuning following SOM inactivation, which depended on different adaptation levels— sharpening the tuning for non-adapted responses but a stronger scaling down effect for adapted responses^55^. To some extent, this may correspond to the two types of modulations observed in our study, while the sharpening effect we found aligns with another recent study using quiet awake mice^54^. Importantly, here we examine the relationship between SOM modulation and sensory discrimination during a behavioral task that demands such processing. We found that type I neurons (tuning curve sharpened by SOM interneurons) also exhibited significant changes in the discrimination of more similar stimuli (**Figure 5E**), suggesting a contribution of SOM interneurons to finer scale frequency discrimination. The mechanisms by which SOM interneurons contribute to the sharpening of tuning curves and the enhancement of stimulus discriminability remain to be investigated. It is conceivable that SOM interneurons might exert feedforward inhibition on responses to non-preferred sensory stimuli and suppress general activity arising from non-sensory inputs, including local cortical recurrent inputs or feedback input from distant brain regions. Distinguishing between these possibilities would be an important topic for future studies.

We only found the scaling effect in a small proportion of neurons during behavior presumably due to differences in behavioral state and stimulus presentations compared to previous studies (**Figure 5I**). Since we imaged from different field of views across experimental sessions, one potential possibility is that the variations in viral expression in SOM neurons across different imaging fields could contribute to the heterogeneity in the effects of photoinhibition. To understand the extent to which such potential variations may influence the observed differential sharpening and scaling effects, we examined the number of type I and type II neurons across different experiments. We found that the number of type I neurons showing sharpening effect are consistently higher than type II neurons showing scaling effect across experiments (**Figure S4**), suggesting that the variations in viral expression is unlikely a significant contributing factor to the differential effects on sensory tuning following SOM inactivation. However, the variation in viral expression in SOM interneurons within a local imaging field could in theory also contribute to the heterogeneity in the identified type I and type II neurons, a possibility we cannot yet rule out due to limited throughput of sampling. In line with the enhancement of neuronal-level discrimination by SOM interneurons, inactivation of SOM interneurons specifically reduced the psychometric slopes at the behavioral level, affecting the discrimination of nearby frequencies but not those farther apart (**Figure 6**). This provides direct evidence for the contribution of SOM interneurons to perceptual discrimination and categorization.

Earlier studies employing activation of interneuron subtypes often reported diverse, sometimes inconsistent effects^6,30,36,58^, potentially due to different network states across experiments^58^. It is important to note that direct activation does not recapitulate physiological activity patterns. For inactivation, however, despite certain caveats such as network-level compensatory effects, the results are more likely to reflect the contribution of endogenous activity. Indeed, by employing inactivation of SOM interneurons, we observed a specific impairment in finer-scale frequency discrimination in perceptual function instead of a general reduction in behavioral performance (**Figure 6**). Such a specific behavioral effect is likely attributable to the neuronal level sharpening effect, also unveiled by inactivation (**Figure 5**).

Although SOM interneurons were modulated by choice behavior, the choice information encoded in SOM interneurons might not be directly transmitted to local populations, as photoinhibition of SOM interneurons did not significantly impact the choice coding of local cortical neurons (**Figure S5**), suggesting that choice information in auditory cortical neurons is more likely to originate from other cortical regions^38,43^.

In our current study, we primarily focused on SOM interneurons in L2/3 of the sensory cortex. The further diversity within SOM interneurons and the differential functions of SOM interneurons in various cortical regions represent important directions for future research, as also demonstrated previously^59^. Lastly, given that SOM interneurons preferentially target distal dendrites of cortical principal neurons, investigating how SOM interneurons regulate dendritic integration during well-defined behavior constitutes another important future direction for unraveling the cellular mechanisms of cortical circuit computations.

## Acknowledgments

We thank J. Pan for technical support; L. Cui and Y. Liu for help in developing the behavioral task. L. Deng for help in histology and animal preparation; C. A. Duan for discussion on data analysis; Z.M. Ying for lab administration.

## Author contributions

R.C. and N.L.X. conceived the project and designed the experiments. R.C. performed the experiments and data analysis. Y.Z. performed the chemogenetic, experiments. R.C. and N.L.X. wrote the manuscript.

## STAR★METHODS

### RESOURCE AVAILABILITY

#### Lead contact

Further information and requests for resources and reagents should be directed to and will be fulfilled by the lead contact, N.L.X.

#### Materials availability

This study did not generate new unique reagents.

#### Data and code availability

The data generated in this study to reproduce all the results have been deposited in the Mendeley Data (doi: 10.17632/pkrv4mgsjw.1). All the original behavioral, optogenetic, chemogenetic, electrophysiological, imaging and tracing data and analysis code are archived in the Center for Excellence in Brain Science and Intelligence Technology, Chinese Academy of Sciences.

## METHODS DETAILS

### Mice

Experimental procedures were approved by the Animal Care and Use Committee of the CAS Center for Excellence in Brain Science and Intelligence Technology, Chinese Academy of Sciences. SOM-IRES-Cre mice were acquired from Jackson lab (Jax number: 013044). Animal age was 8-9 weeks at surgery, and age 9–10 weeks at the start of behavioral training. Mice were group-housed (< 6 mice/cage) in a 12 h reverse light/dark cycle. All experimental procedures were conducted during the dark phase. Mice had no previous history of any other experiments. Mice were water restricted before the start of behavioral training. Each mouse was weighted daily and the body weight was maintained at no less than 85% of the weight before water restriction. On training days, mice received all their water from behavioral task (∼1 ml). Supplementary water was provided for mice who could not maintain a stable body weight from task-related water intake. On days without behavioral training, mice received 1 ml of water. Although it was shown that C57BL/6J mice exhibited significant hearing loss primarily after 7 months old^60^, their hearing threshold are not significantly different in the frequency range of 8-32 kHz during the age of 8-14 weeks^61^. The ages of the animals used in this study are within this range, and the tone stimuli used in this study are also within this frequency range. While it was shown that some C57BL/6J mice may exhibit earlier onset of hearing threshold increase^62^, the sound pressure level used in the current study (70-75 dB) is still well above the hearing threshold within the range of 8-32 kHz.

### Surgery and virus injection

During surgery and virus injection, mice were anaesthetized with isoflurane (1∼2%). For chronic imaging window implantation, a craniotomy of ∼2 mm in diameter was made over the left auditory cortex, with the dura intact. The injection system is as described previously^39^. A pulled glass pipette (25–30 um O.D. at the tip; Drummond Scientific, Wiretrol II Capillary Microdispenser) was filled with mineral oil, and positioned by a Sutter MP-225 manipulator. A metal plunger was inserted into pipette and controlled by a hydraulic manipulator (Narashige, MO-10) to load and inject viral solutions. For SOM neurons imaging, AAV-hSyn-FLEX-GCaMP6s (AAV 2/9, Shanghai Taitool Bioscience Co.Ltd) was slowly injected into left auditory cortex (-2.6 mm AP, 4.7 mm ML from bregma, 300 µm below the dura) of SOM-IRES-Cre mice (10nl∼20nl per minute, 100-150nl per site, 2-3 injection sites per animal). The imaging window is a double-layered glass window (inner layer, 200-um-thick, 2mm in diameter; outer layer, 220-um-thick, 5mm in diameter) adhered using an ultraviolet cured optical adhesive (Norland Optical Adhesives 61). The double-layered glass window was sealed with dental cement (Jet Repair Acrylic, Lang Dental Manufacturing), with the inner layer glass inserted in the craniotomy and outer layer glass anchored on the skull. A titanium head-post was then attached to the skull using cyanoacrylate glue and dental cement. Mice were allowed for at least 7 days to recover before water restriction.

### Head-fixed auditory discrimination behavior

The behavioral apparatus is as described previously^38,39^. Mice were head-fixed and placed in a custom-made double-walled sound-proof box. Mice were head-fixed, with water reward provided by two custom-made metal water spouts placed in front of the mice. The spouts were connected to a capacitive-sensing board that detects the contact of the tongue during licking. Mouse behavior was controlled by the PX-Behavior, a custom-developed real-time system, with the hardware controlled by a microcontroller (Arduino MEGA 2560) and software controlling task protocol and data acquisition written in C++ and python. Sound waveforms were generated from a custom-designed tone-generating module (TGM), sent to an amplifier (ZB1PS, Tucker-Davis Technologies), and delivered through electrostatic speakers (E2, Tucker-Davis Technologies) placed on the right side of mice. The sound systems were calibrated using a free-field microphone (Type 4939, Brüel and Kjær) over 3– 60 kHz and showed a smooth spectrum (± 5 dB). 5 ms cosine ramps were applied to the rise and fall of all tones. The sound intensity was 70-75dB SPL for all tone stimuli.

After water restriction for 3-5 days, during which ∼0.5 ml water per day was provided, mice showed motivation to lick to the spouts to obtain water reward. Behavioral training was started with an initial shaping session, where water was delivered upon licking at either of the lick spouts, alternating every 3 trials. Mice learned to lick left and right in this shaping session. The two-alternative-forced-choice (2AFC) task training was started 2-3 days after initial shaping. In the initial 2AFC task training, mice learned to discriminate two pure tones with 2 octaves apart (8 and 32 kHz, 7 and 28 kHz, or 5 and 20 kHz). Mice were required to lick the left spout following the lower frequency tone and lick the right spout following the higher frequency tone. Each trial was started with a random delay of 500 ms to 2000 ms before sound stimulus onset. The sound stimulus lasts for 300 ms. The answer period (3 s) begins ∼ 500 ms after the offset of sound stimulus. Mice were required to respond by licking within the answer period, during which licking on the correct side would lead to a water reward (∼6 *μ*l), while licking on the wrong side would lead to a time-out punishment of 2-4 s, during which continued licking on the wrong side would reinitiate the time-out period. If mice made no choice in the answer period, the trial was defined as a miss trial. The intertrial interval was 3 s between the end of the current trial and the start of the next trial. Mice with performance >85% (correct rate) for the two easy stimuli (8 kHz and 32 kHz) were included in analysis. It normally took 5-10 training sessions for mice to reach this performance criterion. After mice reached this performance criterion, 4-6 additional tones at intermediate frequencies equally spaced in octaves between the two training tones were delivered to test the categorization performance and sensory tuning of imaged neurons. In the simultaneously manipulating SOM neurons and imaging experiment, besides the six frequencies with equal 0.4 octave distance, there was a tone of frequency at the defined boundary, randomly rewarded on both sides.

We fitted the behavior data with a sigmoid function to obtain the psychometric function of frequency discrimination^39,63,64^

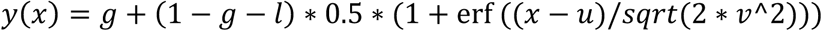

in which *y*(*x*) represents the probability of rightward choice, *x* is octave distance of tone frequencies from the minimal frequency. The parameters: *g* is the guess rate, *l* is the lapse rate, *u* is subject bias, *v* is discrimination threshold (slope in our case). erf () is error function.

### Two photon calcium imaging

We use a custom built two-photon microscope coupled to a Ti-Sapphire laser (Chameleon Ultra II, Coherent) to image the calcium signals in the auditory cortex. The laser was tuned to 920 nm. Signals were collected through a 16 x 0.8 NA objective (Nikon), isolated using a 594 nm high-pass dichroic and a bandpass filter (525/50, Semrock), and detected by GaAsP photomultiplier tubes (10770PB-40, Hamamatsu). Data were acquired using ScanImage at scanning rate of 55 Hz for 256×256 pixels, or 28 Hz for 512 × 512 pixels. The field of view is 447×447 *μ*m for SOM neuron imaging, and 300×300 *μ*m for pyramidal neuron imaging. At the beginning of each trial, the ScanImage receives a trigger from the behavior system to start data acquisition. Imaging experiments were conducted in a double-walled sound-proof box enclosing the entire microscope to eliminate the ambient noise. The high frequency noise from the resonant scanner was blocked using a glass window sealing the metal housing of the scanner. Calcium imaging under passive condition was carried out after mice finished the behavioral task and with the lick ports removed. We acquired imaging data from 846 SOM interneurons from 9 SOM-Cre mice, 30 imaging/behavior sessions. In each session we acquired imaging data from one field of view with identified SOM interneurons. On average, we were able to image from 28±11 SOM interneurons in each field of view.

### Chronic imaging during learning

For imaging experiments across learning, the same imaging field was identified on consecutive days using coarse alignment based on superficial blood vessels followed by careful visual alignment to reference images. Tracking of the same neurons across days was based on visual identification of individual neurons and their surrounding features in zoomed-in images. Only neurons showing clear and similar cell body morphology across sessions were included for analysis. Within the same imaging field, the x-y shift of identified ROIs was manually corrected by moving previous drawn ROIs. All tones of six frequencies were given to mice both in early training stage and expert stage.

### Simultaneous optical inhibition and imaging

AAV-hSyn-GCaMP6s (AAV 2/9) and AAV-hSyn-FLEX-Jaws-KGC-tdTomato-ER2 (AAV 2/9) were mixed with the titer ratio 1:10, and injected in the auditory cortex of SOM-Cre mice to allow the expression of Jaws in SOM interneurons and GCaMP6s in general L2/3 cortical neurons. To inactivate SOM neurons, 635 nm laser light was delivered through the objective via a custom-designed light path with an extra dichroic mirror (FF705-Di01, Semrock) and a modified primary dichroic mirror (FF594-Di04, Semrock) before the entrance pupil of the objective to pass the 635-nm laser into the objective and reflect the GCaMP6s emission light to the detection arm. The 635 nm light coming out of the objective was ∼1 mm in diameter at the focal plan, with the total power of ∼15 mW. The red laser covered 1.5 s after sound onset in photostimulation trials, which were randomly interleaved with control trials at about equal proportions.

### Imaging data analysis

#### Data pre-processing

In initial imaging data analysis, brain motion correction and regions of interest (ROIs) extraction followed the previous study^39^. For SOM neurons, the baseline fluorescence(*F_0_*) of each trial was calculated by averaging the raw fluorescence of the time window (with a duration of 1∼1.5 s) before sound. Then *ΔF/F_0_* was calculated as (*F – F_0_*)/*F_0_*×100%, used as normalized fluorescence change across trials. For pyramidal neurons, imaging was done with continuous scanning, without gap during intertrial interval. Slow calcium fluorescence changes were removed by determining the distribution of fluorescence values in a ∼20 s interval around each sample time point and subtracting the 8th percentile value. For each ROI, *ΔF/F_0_* (%) was calculated as Δ*F/F_0_* × 100, where *F_0_* is the mode of the distribution of *F* obtained from the histogram plot of *F*.

For further analysis of responses under task and passive condition, we used the peak of smoothed activity (by every 5 frames) in a fixed time window after sound. The fixed time window is 1.5 s for SOM neurons, and 1 s for pyramidal neurons, since the onset of calcium signal is slower in SOM neurons. We defined responsive neurons in task using the following criteria. For the preferred tone frequency, at least 70% of trials showed responses exceeding 2 × standard deviations of the pre-stimulus activity (baseline). For comparison of SOM interneurons activity under task with passive conditions, only correct trials were used, to exclude the responses to the opposite categorical choice in the error trial of the same stimulus. In other cases, both correct trials and error trials were used.

#### ROC analysis

To quantify the discrimination for stimuli, categories or choices by single neuron activity, we used an ideal observer decoding based on receiver operating characteristic (ROC) analysis as described previously^43^. The peak during the fixed window (stated above) after stimulus onset were used to construct the ROC curve. The area under ROC curve (auROC) was used as the discriminability of neuronal activity.

#### Generalized linear regression

We built a generalized linear regression model based on 4 behavioral variables: stimulus frequency, categorical choice, reward and lick number

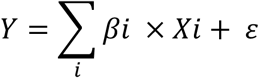

in which *Xi* indicates the *i*th variable. All variables except lick number were defined as categorical variables, *βi* is the regression coefficient of the *i*th variable, *ɛ* represents random noise. We measured the goodness of fit (R^2^) as the variance explained by the model. Neurons with variance explained (R^2^) by full model >0.15 from sessions with >= 5 errors trials in each stimulus category are used for further analysis. The relative contribution of each behavioral variable was measured by training partial models excluding the corresponding variable, calculated as 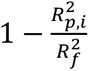, in which 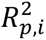 is the R of the partial model excluding the *i*th variable. 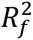 is the R^2^ of the full model. Negative relative contribution was set to zero (in case that R^2^ of the partial model is higher than the full model due to introducing noise).

#### Simultaneously optical manipulation and imaging data processing

In the simultaneously optical manipulation and imaging experiment, the 635 nm red laser illumination led to a small uniform fluorescence increase in the imaging field. To remove this optical artifact, we first quantified the artifact by calculating the mean fluorescence value of non-ROI pixels near each ROI and averaged the time series of this fluorescence value across optical stimulation trials in each session. We then subtracted this averaged optical artifact trace from the fluorescence signal traces for each ROI (Figure S3).

To quantify the effect of optogenetic inactivation of SOM interneurons, we used two-way analysis of variance (ANOVA, using MATLAB function ‘anovan’, linear model), with optical manipulation and frequency as the two contributing factors, to test the significance of each factor contributing to the activity after sound onset of each trial. For each neuron, if p value of the photostimulation factor is below 0.05, and in further multi comparison, p value of at least one frequency is below 0.05, this neuron is defined as significantly modulated by photoinactivation. Comparing the mean activity of all trials in control trials versus photostimulation trials determined the neuron to be significantly increased or decreased.

For pyramidal neurons, we then classified neuronal responses as sensory or choice selective among the responsive neurons (responsive neurons were defined as stated in the ‘Data pre-processing’ section). Choice selective neurons were classified based on 1) two-way ANOVA with frequency and choice as the two contributing factors with the significance level of 0.01 for choice, and 2) fitting the frequency-response relation to a sigmoid function,

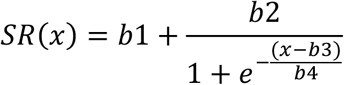

with R^2^ > 0.8. Other responsive neurons which do not fulfill these criteria are classified as sound-tuned neurons.

To characterize neurons sharpened or scaled by SOM interneurons, we calculated the activity difference of each frequency under photostimulation and control condition (D_freq_). If at least one D_freq_ of frequencies excluding best frequency (BF) is greater than the D_freq_ of BF, then it indicates that SOM interneurons exerts more inhibition on other frequency than on BF and we characterized this as neurons sharpened by SOM interneurons (type I). On the contrary, if the activity difference of BF is the largest, it indicates this neuron is inhibited by SOM interneurons in a divisive manner, thus characterized as neurons scaled by SOM interneurons (type II). Tuning of type I neurons under control and photostimulation conditions were then aligned to the best frequency (BF) of control conditions, and normalized by subtracting the activity of the minimal response in each corresponding condition, and divided by the difference of BF response and the minimal response. The normalized responses were fitted with a Gaussian function (**Figure 5C**). Activity of each frequency of type II neurons in photostimulation and control conditions were summarized by subtracting the activity of the minimal response in control condition, and divided by the difference of maximal response and the minimal response in control condition (**Figure 5F**).

Center of mass of tuning curves (COM_tuning_) was calculated as

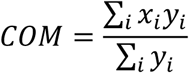

in which *x*_*i*_ is the distance (octave) of the *i*th tone frequency from the lowest frequency, and *y*_*i*_ is the response to the *i*th tone frequency.

### Chemogenetics inhibition

AAV-hSyn-FLEX-hM4Di-mCherry (AAV 2/9) was injected bilaterally into mice auditory cortex. Clozapine N-oxide (CNO, 2-3mg/Kg, in 0.1ml solution) was intraperitoneally (i.p.) injected prior to the experimental sessions. In control sessions, 0.1ml saline was injected. Saline sessions and CNO sessions were interleaved, for paired comparison. Six probe middle frequencies were inserted with the total fraction of 40% trials. To control for the effect of CNO injection only, we injected CNO in mice expressing tdTomato in ACx SOM interneurons (using AAV-hSyn-FLEX-tdTomato-WPRE-bGHpA), and obtained behavioral data.

### Histology

After imaging or manipulation experiments, mice were deeply anesthetized with Chloral hydrate, and perfused with saline and then 0.4% Paraformaldehyde, to get the fixed brain tissue. Brains were fixed with Paraformaldehyde overnight or longer, and then dehydrated with 30% sucrose for at least one day. Then we cut brain slices using a cryostats (Leica CM1950). Images were taken using Olympus VS120.

### Statistics

All data were presented as mean ± SEM, unless mentioned otherwise. Statistics were done using MATLAB R2017a. Statistical significance was tested using paired-sample t test, two-way-ANOVA, Wilcoxon signed-rank test. (ns, p > 0.05, *p < 0.05, **p < 0.01, ***p < 0.001.)

## Supplemental Information

**Figure S1.**
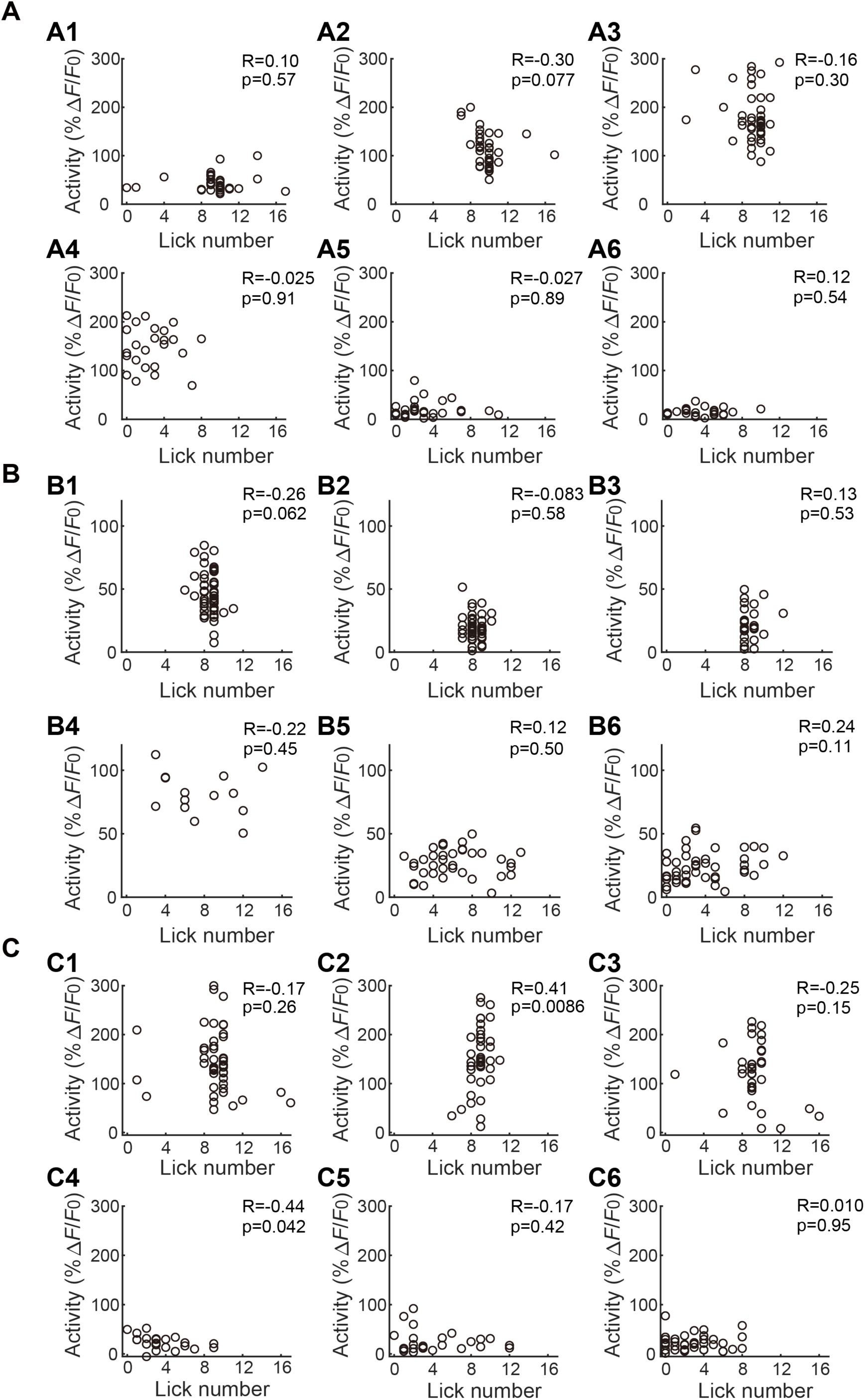
SOM interneuron activity in task is not correlated with lick number. (A)-(C) Correlation analysis of lick number and activity of the 3 example neurons in Figure 1D. Each circle is one trial. A1-A6, B1-B6, C1-C6 correspond to 6 frequencies. The licks during the period from 1 s before to 1.5 s after stimulus onset were used. Neuron activity used is the peak response during the time window of 1.5 s after stimulus onset.

**Figure S2.**
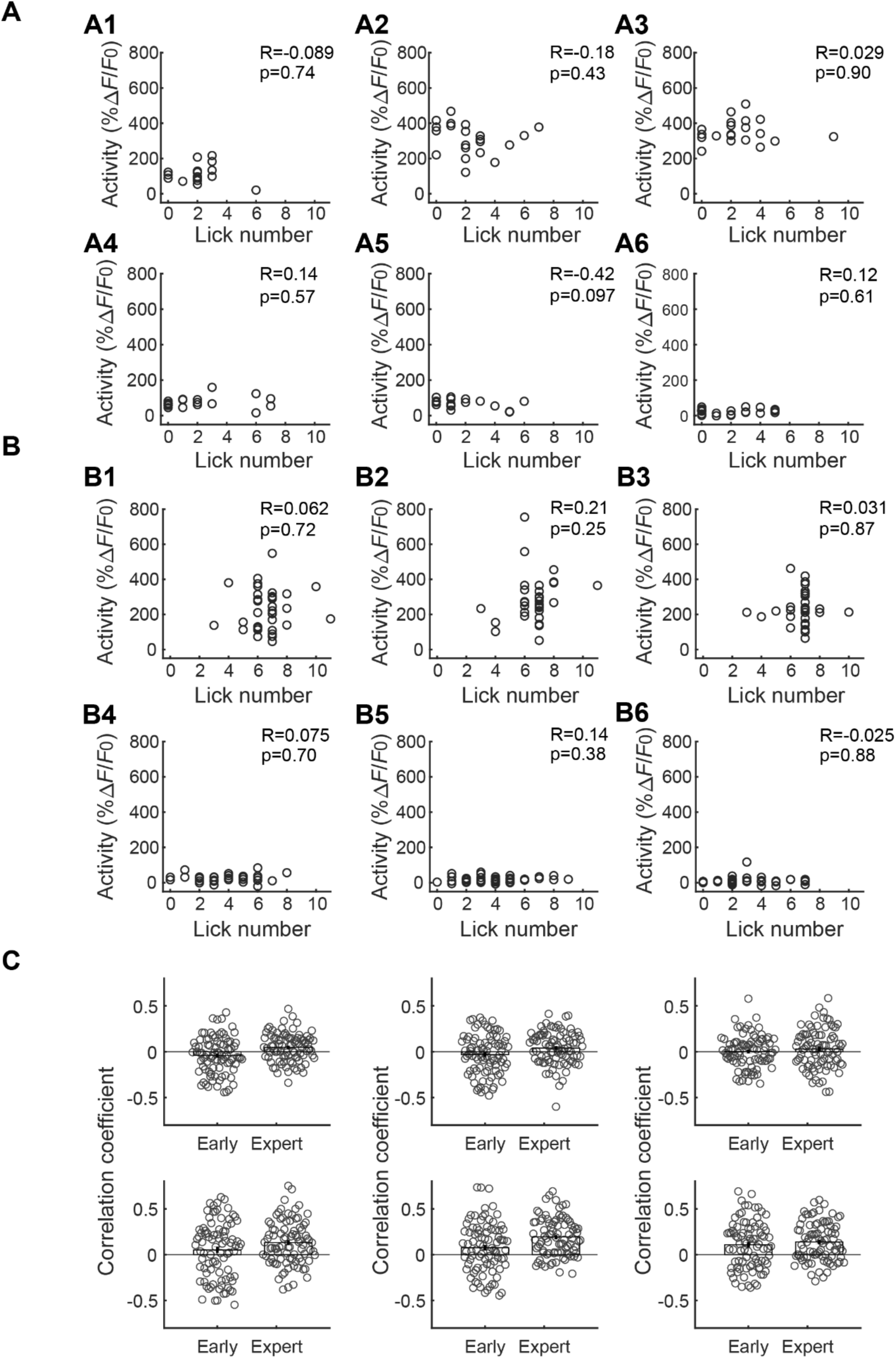
SOM interneuron activity in task is not correlated with lick number across learning. (A) Correlation analysis of lick number and activity of the example neuron in Figure 3A-D, in early learning stage. Each circle is one trial. A1-A6 correspond to 6 frequencies. The licks during the period from 1 s before to 1.5 s after stimulus onset were used. Neuron activity used is the peak response during the time window of 1.5 s after stimulus onset. (B) Same as (A), but in expert learning stage. (C) Summary of correlation analysis of all neurons, showing the coefficients in early and expert stage (n = 88 neurons from 5 imaging fields, 5 mice). Each circle is one neuron. Error bars indicate SEM. 6 panels correspond to 6 frequencies, as illustrated in (A).

**Figure S3.**
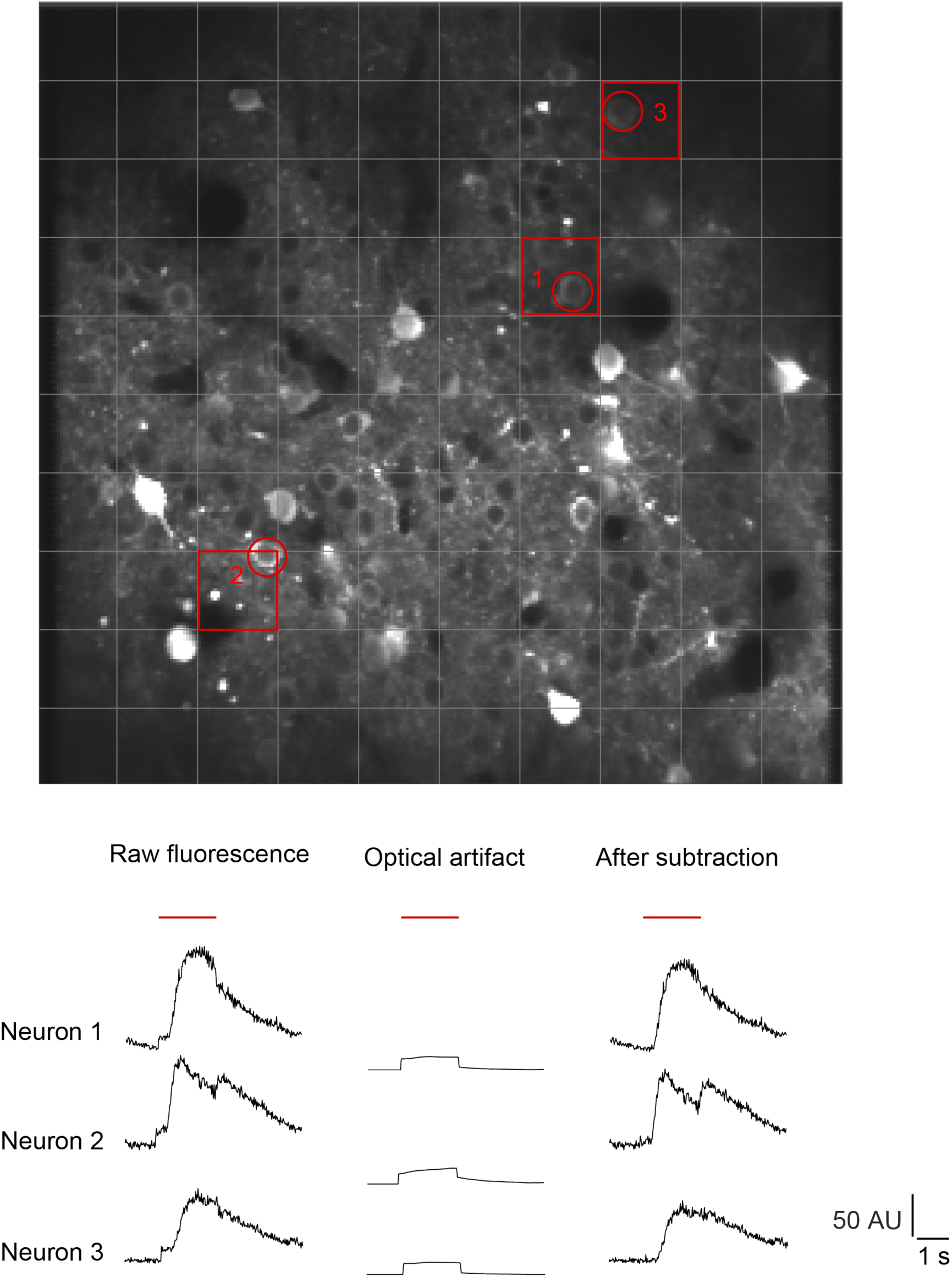
Demonstration of optical artifact removement in simultaneous optogenetic manipulation and two-photon imaging experiments data analysis. Top: an example imaging field with 10 × 10 subregions indicated by the gray grid. We calculated the mean fluorescence value of non-ROI pixels near each ROI in the subregion in which the ROI is located, and averaged the time series of this fluorescence value across optical stimulation trials in each session. We then subtracted this averaged optical artifact trace from the fluorescence signal traces for each ROI. Bottom: three examples showing the effect of artifact subtraction. The 3 neurons and the corresponding subregions are indicated by red circles and red squares in the top image. The red horizontal line indicates laser stimulation.

**Figure S4.**
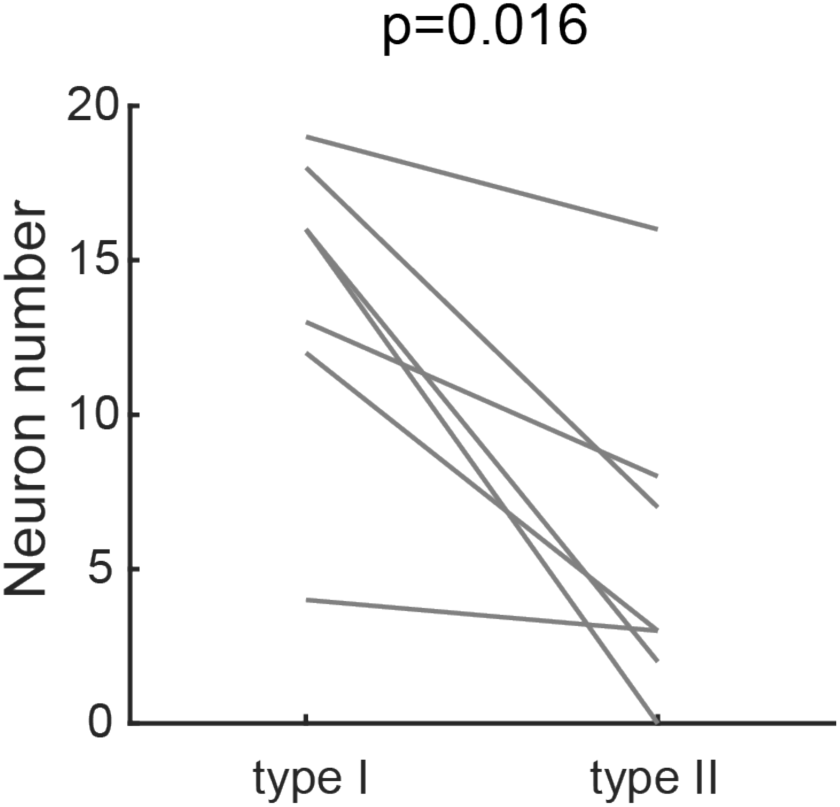
Numbers of type I and type II neurons in each imaging session. Each line represents the numbers of the two types of neurons in the same imaging session (Wilcoxon signed-rank test, n = 7 sessions, 3 mice).

**Figure S5.**
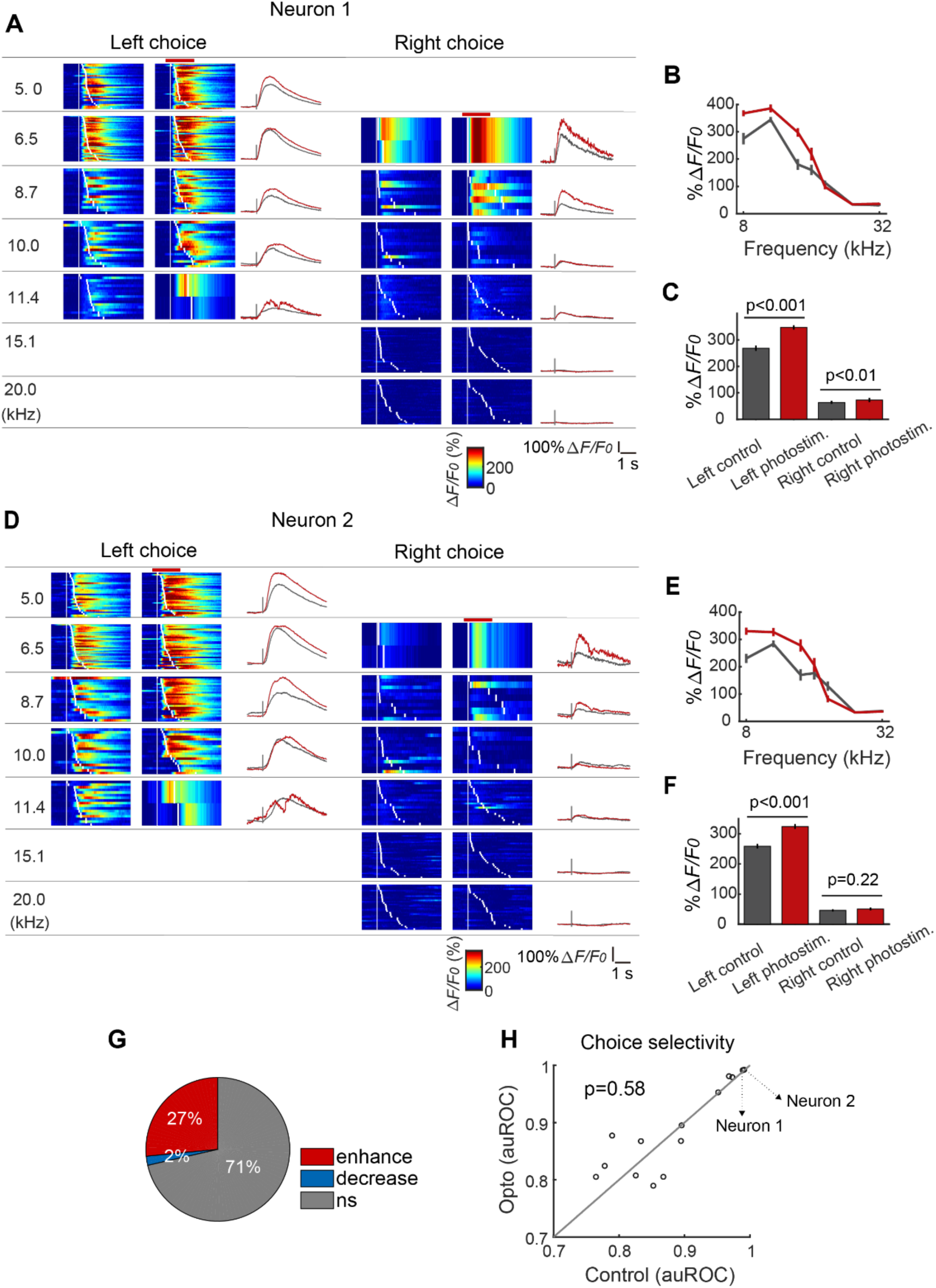
Effect of SOM interneurons inactivation on choice selective neurons. (A) Activity of one example neuron showing choice selectivity. Color raster plots: activity grouped by different tone frequencies, different conditions (control condition and photostimulation condition), and different choices. Activity in each trial is aligned to sound stimulus and laser onset time, indicated by the first white line, and sorted by the time of the first lick after sound onset, indicated by the second white line. The red horizontal line indicates laser stimulation. The traces on the right are the averaged responses across trials in each frequency group. Black, control condition; red, photostimulation condition. The gray line indicates sound stimulus and laser onset time. (B) Responses amplitude for the neuron in (A) as a function of tone frequencies. Black, control condition; red, photostimulation condition. (C) Comparison (using ANOVA) of responses amplitude under control condition and photostimulation condition, within left choice trials and right choice trials. (D) – (F) the same as (A) - (C), for another neuron. (G) Quantification of activity change induced by photostimulation manipulation of all choice selective neurons using ANOVA (n = 49 neurons, from 7 sessions, 3 mice). (H) Choice selectivity (ROC analysis) comparison of control and photostimulation conditions, for all choice selective neurons with significant activity amplitude change by photostimulation manipulation (Wilcoxon signed-rank test, n = 14 neurons, from 7 sessions, 3 mice).

**Figure S6.**
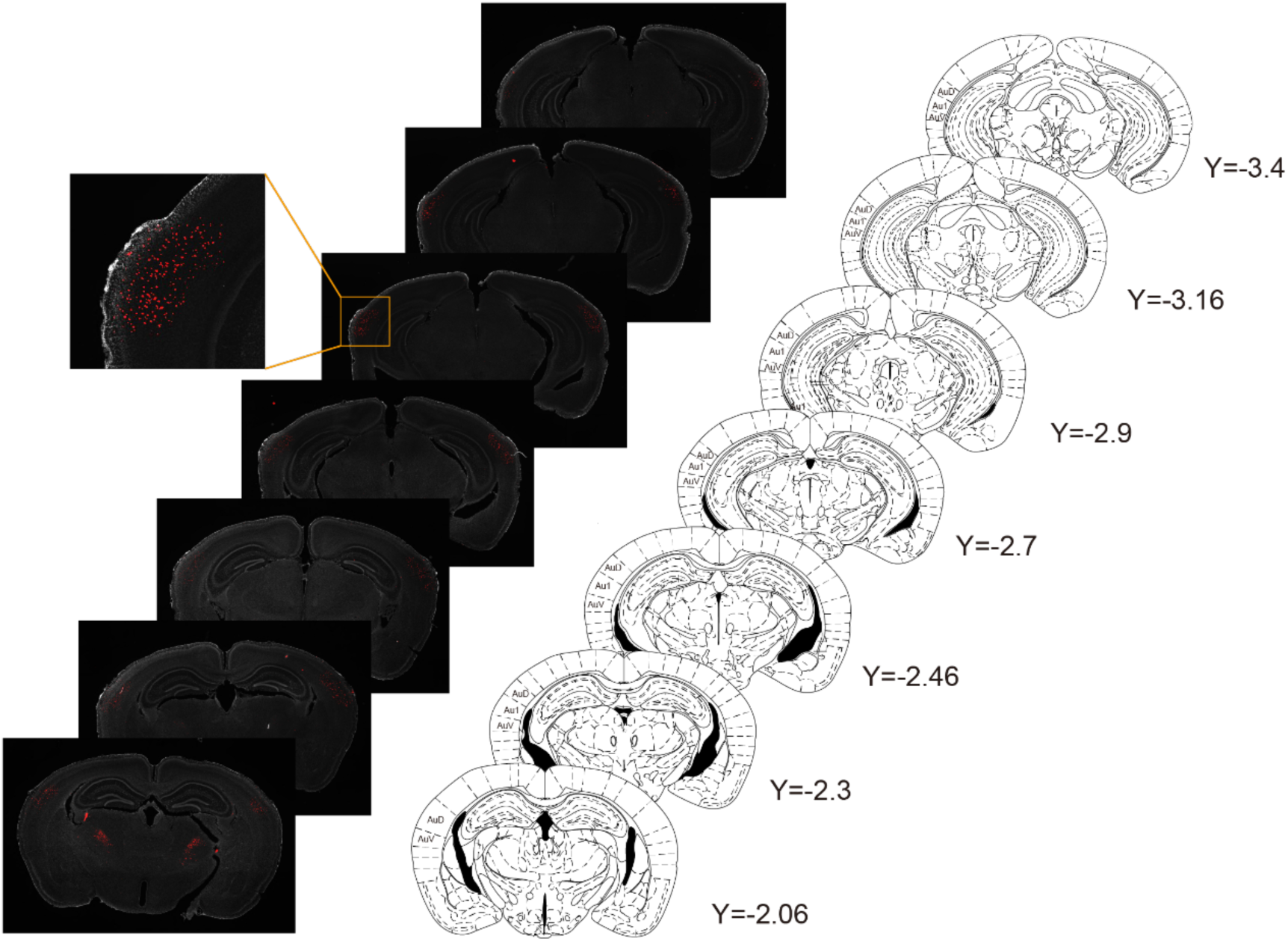
Virus expression range in the SOM interneurons chemogenetic manipulation experiment. Brain slice histology images of one example animal expressing hM4Di-mCherry in auditory cortex, arranged in anterior-posterior axis order, in reference to the brain atlas. For visualization, the image is set as binary. Signals above the threshold can be visualized, whereas others cannot. Then the image is merged with DAPI channel, displaying in gray. Zoomed image of the region in the square is shown. The red signal in regions far from injection site is confirmed to be autofluorescence.

**Table S1.** Number of neurons across imaging fields for SOM interneurons imaging.

| Animal No. | imaging field No. | Neuron number |
| --- | --- | --- |
| 1 | 1 | 20 |
| 2 | 1 | 19 |
|  | 2 | 18 |
| 3 | 1 | 32 |
|  | 2 | 27 |
|  | 3 | 58 |
|  | 4 | 41 |
| 4 | 1 | 30 |
|  | 2 | 34 |
|  | 3 | 37 |
|  | 4 | 61 |
| 5 | 1 | 10 |
| 6 | 1 | 7 |
| 7 | 1 | 33 |
|  | 2 | 27 |
|  | 3 | 26 |
|  | 4 | 33 |
|  | 5 | 21 |
| 8 | 1 | 19 |
|  | 2 | 33 |
|  | 3 | 36 |
|  | 4 | 31 |
|  | 5 | 32 |
|  | 6 | 15 |
| 9 | 1 | 16 |
|  | 2 | 27 |
|  | 3 | 19 |
|  | 4 | 23 |
|  | 5 | 33 |
|  | 6 | 28 |

**Table S2.** Number of neurons across imaging fields for SOM interneuron activity imaging across learning.

| Animal No. | imaging field No. | Neuron number |
| --- | --- | --- |
| 10 | 1 | 13 |
| 11 | 1 | 14 |
| 12 | 1 | 22 |
| 13 | 1 | 16 |
| 14 | 1 | 23 |

**Table S3.** Number of neurons across imaging fields for simultaneous optogenetic manipulation of SOM interneurons and imaging in auditory cortex.

| Animal No. | imaging field No. | Neuron number |
| --- | --- | --- |
| 15 | 1 | 100 (4) |
|  | 2 | 85 (5) |
| 16 | 1 | 87 (3) |
|  | 2 | 104 (4) |
|  | 3 | 101 (3) |
| 17 | 1 | 101 (5) |
|  | 2 | 123 (2) |

Note: the neuron numbers correspond to putatively pyramidal neurons, and the numbers inside the parentheses correspond to SOM interneuron numbers in the same field.

## References

1. Fishell, G., and Rudy, B. (2011). Mechanisms of Inhibition within the Telencephalon: “Where the Wild Things Are.” Annu. Rev. Neurosci. 34, 535–567. 10.1146/annurev-neuro-061010-113717.

2. Kepecs, A., and Fishell, G. (2014). Interneuron cell types are fit to function. Nature 505, 318–326. 10.1038/nature12983.

3. Markram, H., Toledo-Rodriguez, M., Wang, Y., Gupta, A., Silberberg, G., and Wu, C. (2004). Interneurons of the neocortical inhibitory system. Nature Reviews Neuroscience 5, 793–807. 10.1038/nrn1519.

4. Tremblay, R., Lee, S., and Rudy, B. (2016). GABAergic Interneurons in the Neocortex: From Cellular Properties to Circuits. Neuron 91, 260–292. 10.1016/j.neuron.2016.06.033.

5. Iaccarino, H.F., Singer, A.C., Martorell, A.J., Rudenko, A., Gao, F., Gillingham, T.Z., Mathys, H., Seo, J., Kritskiy, O., Abdurrob, F., et al. (2016). Gamma frequency entrainment attenuates amyloid load and modifies microglia. Nature 540, 230–235. 10.1038/nature20587.

6. Lee, S.-H., Kwan, A.C., Zhang, S., Phoumthipphavong, V., Flannery, J.G., Masmanidis, S.C., Taniguchi, H., Huang, Z.J., Zhang, F., Boyden, E.S., et al. (2012). Activation of specific interneurons improves V1 feature selectivity and visual perception. Nature 488, 379–383. 10.1038/nature11312.

7. Letzkus, J.J., Wolff, S.B.E., Meyer, E.M.M., Tovote, P., Courtin, J., Herry, C., and Lüthi, A. (2011). A disinhibitory microcircuit for associative fear learning in the auditory cortex. Nature 480, 331–335. 10.1038/nature10674.

8. Schneider, D.M., Nelson, A., and Mooney, R. (2014). A synaptic and circuit basis for corollary discharge in the auditory cortex. Nature 513, 189–194. 10.1038/nature13724.

9. Sohal, V.S., Zhang, F., Yizhar, O., and Deisseroth, K. (2009). Parvalbumin neurons and gamma rhythms enhance cortical circuit performance. Nature 459, 698–702. 10.1038/nature07991.

10. Yizhar, O., Fenno, L.E., Prigge, M., Schneider, F., Davidson, T.J., O’Shea, D.J., Sohal, V.S., Goshen, I., Finkelstein, J., Paz, J.T., et al. (2011). Neocortical excitation/inhibition balance in information processing and social dysfunction. Nature 477, 171–178. 10.1038/nature10360.

11. Fu, Y., Tucciarone, J.M., Espinosa, J.S., Sheng, N., Darcy, D.P., Nicoll, R.A., Huang, Z.J., and Stryker, M.P. (2014). A Cortical Circuit for Gain Control by Behavioral State. Cell 156, 1139–1152. 10.1016/j.cell.2014.01.050.

12. Lee, S., Kruglikov, I., Huang, Z.J., Fishell, G., and Rudy, B. (2013). A disinhibitory circuit mediates motor integration in the somatosensory cortex. Nat Neurosci 16, 1662–1670. 10.1038/nn.3544.

13. Pi, H.-J., Hangya, B., Kvitsiani, D., Sanders, J.I., Huang, Z.J., and Kepecs, A. (2013). Cortical interneurons that specialize in disinhibitory control. Nature 503, 521–524. 10.1038/nature12676.

14. Urban-Ciecko, J., and Barth, A.L. (2016). Somatostatin-expressing neurons in cortical networks. Nat Rev Neurosci 17, 401–409. 10.1038/nrn.2016.53.

15. Chiu, C.Q., Lur, G., Morse, T.M., Carnevale, N.T., Ellis-Davies, G., and Higley, M.J. (2013). Compartmentalization of GABAergic Inhibition by Dendritic Spines. Science 340, 759–762. 10.1126/science.1234274.

16. Murayama, M., Pérez-Garci, E., Nevian, T., Bock, T., Senn, W., and Larkum, M.E. (2009). Dendritic encoding of sensory stimuli controlled by deep cortical interneurons. Nature 457, 1137–1141. 10.1038/nature07663.

17. Silberberg, G., and Markram, H. (2007). Disynaptic inhibition between neocortical pyramidal cells mediated by Martinotti cells. Neuron 53, 735–746. 10.1016/j.neuron.2007.02.012.

18. Muñoz, W., Tremblay, R., Levenstein, D., and Rudy, B. (2017). Layer-specific modulation of neocortical dendritic inhibition during active wakefulness. Science 355, 954–959. 10.1126/science.aag2599.

19. Cottam, J.C.H., Smith, S.L., and Häusser, M. (2013). Target-Specific Effects of Somatostatin-Expressing Interneurons on Neocortical Visual Processing. J Neurosci 33, 19567–19578. 10.1523/JNEUROSCI.2624-13.2013.

20. Pfeffer, C.K., Xue, M., He, M., Huang, Z.J., and Scanziani, M. (2013). Inhibition of inhibition in visual cortex: the logic of connections between molecularly distinct interneurons. Nat Neurosci 16, 1068–1076. 10.1038/nn.3446.

21. Liu, D., Deng, J., Zhang, Z., Zhang, Z.-Y., Sun, Y.-G., Yang, T., and Yao, H. (2020). Orbitofrontal control of visual cortex gain promotes visual associative learning. Nat Commun 11, 2784. 10.1038/s41467-020-16609-7.

22. Chen, N., Sugihara, H., and Sur, M. (2015). An acetylcholine-activated microcircuit drives temporal dynamics of cortical activity. Nat Neurosci advance online publication. 10.1038/nn.4002.

23. Kato, H.K., Gillet, S.N., and Isaacson, J.S. (2015). Flexible Sensory Representations in Auditory Cortex Driven by Behavioral Relevance. Neuron 88, 1027–1039. 10.1016/j.neuron.2015.10.024.

24. Khan, A.G., Poort, J., Chadwick, A., Blot, A., Sahani, M., Mrsic-Flogel, T.D., and Hofer, S.B. (2018). Distinct learning-induced changes in stimulus selectivity and interactions of GABAergic interneuron classes in visual cortex. Nature Neuroscience 21, 851–859. 10.1038/s41593-018-0143-z.

25. Kuchibhotla, K.V., Gill, J.V., Lindsay, G.W., Papadoyannis, E.S., Field, R.E., Hindmarsh Sten, T.A., Miller, K.D., and Froemke, R.C. (2017). Parallel processing by cortical inhibition enables context-dependent behavior. Nat Neurosci 20, 62–71. 10.1038/nn.4436.

26. Pakan, J.M., Lowe, S.C., Dylda, E., Keemink, S.W., Currie, S.P., Coutts, C.A., and Rochefort, N.L. (2016). Behavioral-state modulation of inhibition is context-dependent and cell type specific in mouse visual cortex. eLife 5, e14985. 10.7554/eLife.14985.

27. Kato, H.K., Asinof, S.K., and Isaacson, J.S. (2017). Network-Level Control of Frequency Tuning in Auditory Cortex. Neuron 95, 412–423.e4. 10.1016/j.neuron.2017.06.019.

28. Chen, G., Zhang, Y., Li, X., Zhao, X., Ye, Q., Lin, Y., Tao, H.W., Rasch, M.J., and Zhang, X. (2017). Distinct Inhibitory Circuits Orchestrate Cortical beta and gamma Band Oscillations. Neuron 96, 1403–1418.e6. 10.1016/j.neuron.2017.11.033.

29. Veit, J., Hakim, R., Jadi, M.P., Sejnowski, T.J., and Adesnik, H. (2017). Cortical gamma band synchronization through somatostatin interneurons. Nature Neuroscience 20, 951–959. 10.1038/nn.4562.

30. Atallah, B.V., Bruns, W., Carandini, M., and Scanziani, M. (2012). Parvalbumin-Expressing Interneurons Linearly Transform Cortical Responses to Visual Stimuli. Neuron 73, 159–170. 10.1016/j.neuron.2011.12.013.

31. Atallah, B.V., Scanziani, M., and Carandini, M. (2014). Atallah et al. reply. Nature 508, E3–E3. 10.1038/nature13129.

32. El-Boustani, S., Wilson, N.R., Runyan, C.A., and Sur, M. (2014). El-Boustani et al. reply. Nature 508, E3–E4. 10.1038/nature13130.

33. El-Boustani, S., and Sur, M. (2014). Response-dependent dynamics of cell-specific inhibition in cortical networks in vivo. Nat Commun 5, 5689. 10.1038/ncomms6689.

34. Lee, S.-H., Kwan, A.C., and Dan, Y. (2014). Interneuron subtypes and orientation tuning. Nature 508, E1–E2. 10.1038/nature13128.

35. Phillips, E.A., and Hasenstaub, A.R. (2016). Asymmetric effects of activating and inactivating cortical interneurons. eLife 5, e18383. 10.7554/eLife.18383.

36. Wilson, N.R., Runyan, C.A., Wang, F.L., and Sur, M. (2012). Division and subtraction by distinct cortical inhibitory networks in vivo. Nature 488, 343–348. 10.1038/nature11347.

37. Weible, A.P., Moore, A.K., Liu, C., DeBlander, L., Wu, H., Kentros, C., and Wehr, M. (2014). Perceptual Gap Detection Is Mediated by Gap Termination Responses in Auditory Cortex. Current Biology 24, 1447–1455. 10.1016/j.cub.2014.05.031.

38. Zhong, L., Zhang, Y., Duan, C.A., Deng, J., Pan, J., and Xu, N. (2019). Causal contributions of parietal cortex to perceptual decision-making during stimulus categorization. Nature Neuroscience 22, 963. 10.1038/s41593-019-0383-6.

39. Xin, Y., Zhong, L., Zhang, Y., Zhou, T., Pan, J., and Xu, N. (2019). Sensory-to-Category Transformation via Dynamic Reorganization of Ensemble Structures in Mouse Auditory Cortex. Neuron 103, 909–921.e6. 10.1016/j.neuron.2019.06.004.

40. Chen, T.-W., Wardill, T.J., Sun, Y., Pulver, S.R., Renninger, S.L., Baohan, A., Schreiter, E.R., Kerr, R.A., Orger, M.B., Jayaraman, V., et al. (2013). Ultrasensitive fluorescent proteins for imaging neuronal activity. Nature 499, 295–300. 10.1038/nature12354.

41. Tang, S., Cui, L., Pan, J., and Xu, N. (2024). Dynamic ensemble balance in direct- and indirect-pathway striatal projection neurons underlying decision-related action selection. Cell Reports 43. 10.1016/j.celrep.2024.114726.

42. Chuong, A.S., Miri, M.L., Busskamp, V., Matthews, G.A.C., Acker, L.C., Sørensen, A.T., Young, A., Klapoetke, N.C., Henninger, M.A., Kodandaramaiah, S.B., et al. (2014). Noninvasive optical inhibition with a red-shifted microbial rhodopsin. Nat Neurosci 17, 1123–1129. 10.1038/nn.3752.

43. Liu, Y., Xin, Y., and Xu, N. (2021). A cortical circuit mechanism for structural knowledge-based flexible sensorimotor decision-making. Neuron 109, 2009–2024.e6. 10.1016/j.neuron.2021.04.014.

44. Yang, Y., Liu, D., Huang, W., Deng, J., Sun, Y., Zuo, Y., and Poo, M. (2016). Selective synaptic remodeling of amygdalocortical connections associated with fear memory. Nat Neurosci 19, 1348–1355. 10.1038/nn.4370.

45. Armbruster, B.N., Li, X., Pausch, M.H., Herlitze, S., and Roth, B.L. (2007). Evolving the lock to fit the key to create a family of G protein-coupled receptors potently activated by an inert ligand. Proc Natl Acad Sci U S A 104, 5163–5168. 10.1073/pnas.0700293104.

46. Cobb, S.R., Buhl, E.H., Halasy, K., Paulsen, O., and Somogyi, P. (1995). Synchronization of neuronal activity in hippocampus by individual GABAergic interneurons. Nature 378, 75–78. 10.1038/378075a0.

47. Kapfer, C., Glickfeld, L.L., Atallah, B.V., and Scanziani, M. (2007). Supralinear increase of recurrent inhibition during sparse activity in the somatosensory cortex. Nat Neurosci 10, 743–753. 10.1038/nn1909.

48. Xu, H., Jeong, H.-Y., Tremblay, R., and Rudy, B. (2013). Neocortical Somatostatin-expressing GABAergic Interneurons Disinhibit the Thalamorecipient Layer 4. Neuron 77, 155–167. 10.1016/j.neuron.2012.11.004.

49. Huang, Z.J., and Zeng, H. (2013). Genetic Approaches to Neural Circuits in the Mouse. Annu. Rev. Neurosci. 36, 183–215. 10.1146/annurev-neuro-062012-170307.

50. Luo, L., Callaway, E.M., and Svoboda, K. (2018). Genetic Dissection of Neural Circuits: A Decade of Progress. Neuron 98, 256–281. 10.1016/j.neuron.2018.03.040.

51. Fenno, L., Yizhar, O., and Deisseroth, K. (2011). The Development and Application of Optogenetics. Annual Review of Neuroscience 34, 389–412. 10.1146/annurev-neuro-061010-113817.

52. Scanziani, M., and Häusser, M. (2009). Electrophysiology in the age of light. Nature 461, 930–939. 10.1038/nature08540.

53. Adesnik, H., and Scanziani, M. (2010). Lateral competition for cortical space by layer-specific horizontal circuits. Nature 464, 1155–1160. 10.1038/nature08935.

54. Lakunina, A.A., Nardoci, M.B., Ahmadian, Y., and Jaramillo, S. (2020). Somatostatin-Expressing Interneurons in the Auditory Cortex Mediate Sustained Suppression by Spectral Surround. J. Neurosci. 40, 3564–3575. 10.1523/JNEUROSCI.1735-19.2020.

55. Natan, R.G., Rao, W., and Geffen, M.N. (2017). Cortical Interneurons Differentially Shape Frequency Tuning following Adaptation. Cell Reports 21, 878–890. 10.1016/j.celrep.2017.10.012.

56. Li, L., Ji, X., Liang, F., Li, Y., Xiao, Z., Tao, H.W., and Zhang, L.I. (2014). A Feedforward Inhibitory Circuit Mediates Lateral Refinement of Sensory Representation in Upper Layer 2/3 of Mouse Primary Auditory Cortex. J. Neurosci. 34, 13670–13683. 10.1523/JNEUROSCI.1516-14.2014.

57. Yu, J., Hu, H., Agmon, A., and Svoboda, K. (2019). Recruitment of GABAergic Interneurons in the Barrel Cortex during Active Tactile Behavior. Neuron 104, 412–427.e4. 10.1016/j.neuron.2019.07.027.

58. Seybold, B.A., Phillips, E.A.K., Schreiner, C.E., and Hasenstaub, A.R. (2015). Inhibitory Actions Unified by Network Integration. Neuron 87, 1181–1192. 10.1016/j.neuron.2015.09.013.

59. Kvitsiani, D., Ranade, S., Hangya, B., Taniguchi, H., Huang, J.Z., and Kepecs, A. (2013). Distinct behavioural and network correlates of two interneuron types in prefrontal cortex. Nature 498, 363–366. 10.1038/nature12176.

60. Hequembourg, S., and Liberman, M.C. (2001). Spiral Ligament Pathology: A Major Aspect of Age-Related Cochlear Degeneration in C57BL/6 Mice. Journal of the Association for Research in Otolaryngology 2, 118–129. 10.1007/s101620010075.

61. Ison, J.R., Agrawal, P., Pak, J., and Vaughn, W.J. (1998). Changes in temporal acuity with age and with hearing impairment in the mouse: a study of the acoustic startle reflex and its inhibition by brief decrements in noise level. J. Acoust. Soc. Am. 104, 1696–1704.

62. Ison, J.R., Allen, P.D., and O’Neill, W.E. (2007). Age-Related Hearing Loss in C57BL/6J Mice has both Frequency-Specific and Non-Frequency-Specific Components that Produce a Hyperacusis-Like Exaggeration of the Acoustic Startle Reflex. JARO 8, 539–550. 10.1007/s10162-007-0098-3.

63. Carandini, M., and Churchland, A.K. (2013). Probing perceptual decisions in rodents. Nat Neurosci 16, 824–831. 10.1038/nn.3410.

64. Wichmann, F.A., and Hill, N.J. (2001). The psychometric function: I. Fitting, sampling, and goodness of fit. Perception & Psychophysics 63, 1293–1313. 10.3758/BF03194544.

